# Structure-Based Design with Tag-Based Purification and In-Process Biotinylation Enable Streamlined Development of SARS-CoV-2 Spike Molecular Probes

**DOI:** 10.1101/2020.06.22.166033

**Authors:** Tongqing Zhou, I-Ting Teng, Adam S. Olia, Gabriele Cerutti, Jason Gorman, Alexandra Nazzari, Wei Shi, Yaroslav Tsybovsky, Lingshu Wang, Shuishu Wang, Baoshan Zhang, Yi Zhang, Phinikoula S. Katsamba, Yuliya Petrova, Bailey B. Banach, Ahmed S. Fahad, Lihong Liu, Sheila N. Lopez Acevedo, Bharat Madan, Matheus Oliveira de Souza, Xiaoli Pan, Pengfei Wang, Jacy R. Wolfe, Michael Yin, David D. Ho, Emily Phung, Anthony DiPiazza, Lauren Chang, Olubukula Abiona, Kizzmekia S. Corbett, Brandon J. DeKosky, Barney S. Graham, John R. Mascola, John Misasi, Tracy Ruckwardt, Nancy J. Sullivan, Lawrence Shapiro, Peter D. Kwong

**Affiliations:** Vaccine Research Center, National Institute of Allergy and Infectious Diseases, National Institutes of Health, Bethesda, MD 20892, USA; Department of Biochemistry and Molecular Biophysics, Columbia University, New York, NY 10032, USA; Electron Microscopy Laboratory, Cancer Research Technology Program, Leidos Biomedical Research Inc., Frederick National Laboratory for Cancer Research, Frederick, MD 21701, USA; Bioengineering Graduate Program, The University of Kansas, Lawrence, KS 66045, USA; Department of Pharmaceutical Chemistry, The University of Kansas, Lawrence, KS 66045, USA; Aaron Diamond AIDS Research Center, Columbia University Vagelos College of Physicians and Surgeons, New York, NY 10032, USA; Department of Chemical Engineering, The University of Kansas, Lawrence, KS 66045, USA

**Keywords:** antibody, biotinylated probe, COVID-19, HRV3C protease, single-chain Fc, structure-based design

## Abstract

Biotin-labeled molecular probes, comprising specific regions of the SARS-CoV-2 spike, would be helpful in the isolation and characterization of antibodies targeting this recently emerged pathogen. To develop such probes, we designed constructs incorporating an N-terminal purification tag, a site-specific protease-cleavage site, the probe region of interest, and a C-terminal sequence targeted by biotin ligase. Probe regions included full-length spike ectodomain as well as various subregions, and we also designed mutants to eliminate recognition of the ACE2 receptor. Yields of biotin-labeled probes from transient transfection ranged from ∼0.5 mg/L for the complete ectodomain to >5 mg/L for several subregions. Probes were characterized for antigenicity and ACE2 recognition, and the structure of the spike ectodomain probe was determined by cryo-electron microscopy. We also characterized antibody-binding specificities and cell-sorting capabilities of the biotinylated probes. Altogether, structure-based design coupled to efficient purification and biotinylation processes can thus enable streamlined development of SARS-CoV-2 spike-ectodomain probes.

## Introduction

Severe acute respiratory syndrome coronavirus 2 (SARS-CoV-2), the causative agent for Coronavirus Disease 2019 (COVID-19), emerged in 2019 and rapidly spread, infecting millions, overwhelming health-care systems, and impacting economies worldwide (Callaway et al., 2020; Cucinotta and Vanelli, 2020). To respond, a global effort has been initiated to develop vaccines and therapeutic agents. For COVID-19 vaccine development (reviewed in Callaway, 2020), the trimeric SARS-CoV-2 spike – a type 1 fusion machine that facilitates virus-cell entry through interaction with the ACE2 receptor (Hoffmann et al., 2020; Ou et al., 2020) – is a lead target, as antibodies against it can block virus entry (Jiang et al., 2020). Most of the SARS-CoV-2 neutralizing antibodies so far isolated target the receptor binding domain (RBD) of the spike protein (Brouwer et al., 2020; Cao et al., 2020; Chen et al., 2020; Chi et al., 2020; Ju et al., 2020; Liu et al., 2020; Pinto et al., 2020; Robbiani et al., 2020; Rogers et al., 2020; Seydoux et al., 2020; Wang et al., 2020a; Wrapp et al., 2020a; Wu et al., 2020; Zeng et al., 2020; Zost et al., 2020), but there are other sites in the N-terminal domain and S2 stem domain that have also been associated with neutralizing activity against other betacoronaviruses (Pallesen et al., 2017; Wang et al., 2018b). Such virus-neutralizing antibodies are sought as therapeutic and prophylactic agents (Cao et al., 2020; reviewed in Graham et al., 2013; Zhou and Zhao, 2020).

Biotin-labeled molecular probes, comprising the SARS-CoV-2 spike as well as its discrete domains, can accelerate development of both vaccines and therapeutic antibodies. For vaccine development, such probes can be used to track humoral responses longitudinally (Liu et al., 2011; Yongchen et al., 2020) and to quantify elicited responses against spike and its domains, as correlating such responses with neutralization should provide crucial insight into sites of spike vulnerability. For antibody identification, probes are used in B cell sorting to identify B cells encoding antibodies capable of recognizing the spike or particular spike domains as well as characterization of antibody binding affinities through surface plasmon resonance (SPR) or bio-layer interferometry (BLI) analyses.

Here we describe the structure-based design of molecular probes, encompassing SARS-CoV-2 spike and its domains. We first designed a construct that allowed for tag-based purification and on-column biotinylation. Next, we incorporated the SARS-CoV-2 spike ectodomain, with prefusion stabilizing mutations and a C-terminal trimerization motif, which we expressed, biotinylated, purified, and characterized, including by cryo-EM. Based on the structure-defined spike-domain organization (Walls et al., 2020; Wrapp et al., 2020b), we also created and characterized separate molecular probes comprising the N-terminal domain (NTD), the receptor-binding domain (RBD), and RBD with spike domain 1 (RBD-SD1). We also used the structure of RBD with ACE2 (Lan et al., 2020; Wang et al., 2020b; Yan et al., 2020a) to define mutations that could inhibit ACE2 interaction, which we incorporated into mutant RBD probes with ACE2-recognition ablated. Finally, we characterized properties of the devised probes including degree of biotinylation, antibody-binding specificity, and use in sorting yeast cells expressing SARS-CoV-2 spike-binding antibodies or B cells from a COVID-19 convalescent donor. Overall, our findings demonstrate how structure-based design can be used to develop molecular probes of the SARS-CoV-2 spike.

## Results

### Strategy for Tag-Based Purification with In-Process Biotinylation

To facilitate purification and biotinylation, we devised a probe construct using the constant portion of an antibody (Fc) as an N-terminal purification tag. Fc expresses and folds efficiently and can be effectively captured by protein A resins (Jager et al., 2013). To prevent intermolecular dimer formation of the Fc-tag, we used a single chain-Fc (scFc) with ‘knob-in-hole’ feature (Ridgway et al., 1996), in which a “knob” comprising a protruding tyrosine (T366Y) was incorporated into the N-terminal-half of the Fc-interface, followed by a 20-residue-linker (GGGGS)_4_ and a “hole” comprising a recessed threonine (Y407T) was incorporated into the C-terminal-half Fc-interface [Kabat antibody residue numbering (Johnson and Wu, 2000)] (Figure 1A).

**Figure 1.**
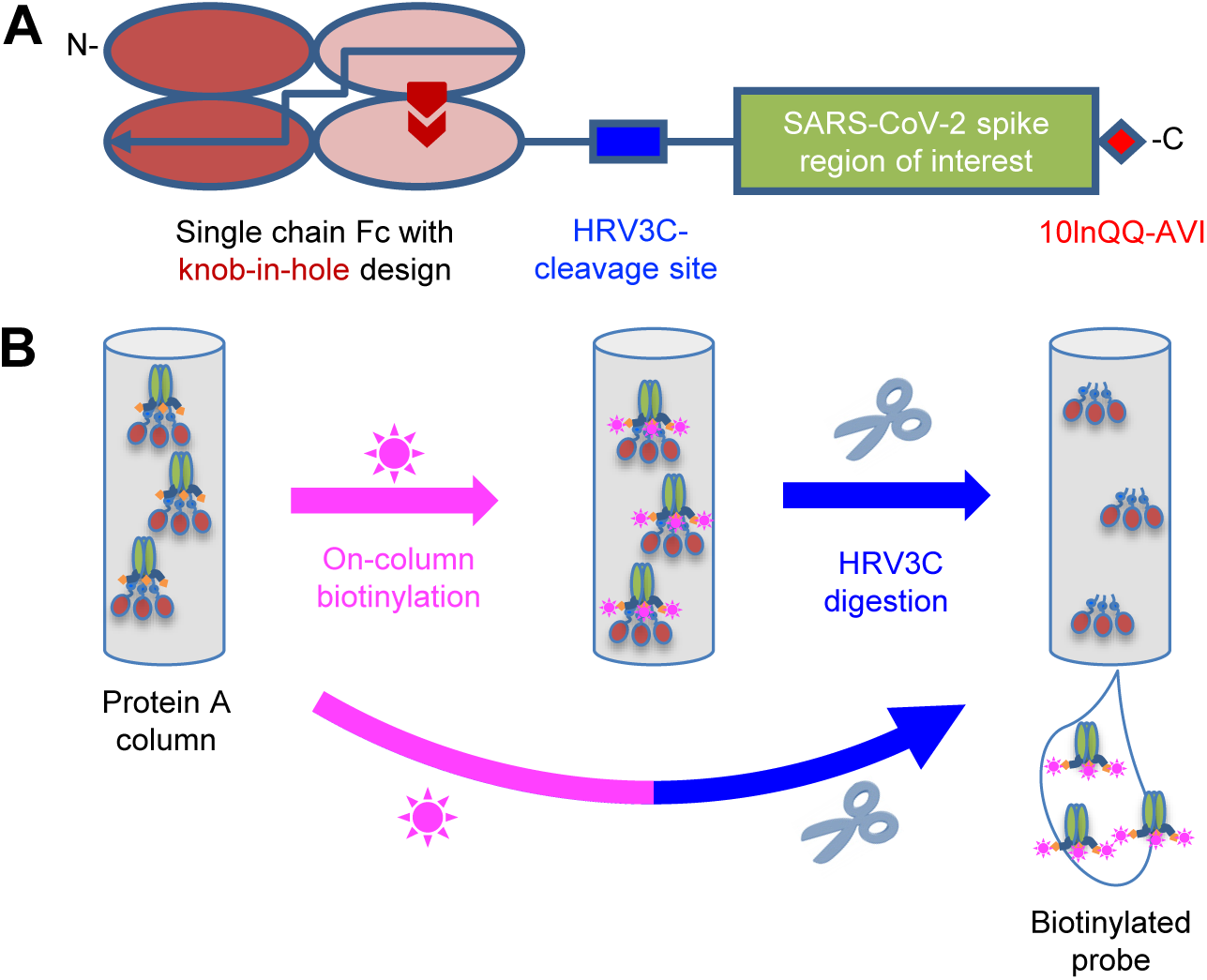
Strategy for Tag-Based Purification with On-Column Biotinylation. (A) Schematic design of the expression construct of SARS-CoV-2 molecular probes. A single chain human Ig constant domain (scFc) was added at N terminus to facilitate expression and purification. The AVI tag was placed at the C terminus after a 10-amino acid linker for biotinylation. The red arrows in the second and fourth Fc domains showed the “knob-in-hole” mutations to prevent dimerization of the scFc. (B) Biotinylation and HRV3C digestion. Cell culture supernatant from cells transiently transfected with plasmid was loaded onto protein A affinity column. Biotinylation and HRV3C cleavage reactions can be carried out in series or simultaneously, as buffers for both reactions are compatible.

Between the scFc and the target region of interest (Figure 1B), we added a cleavage site for the human rhinovirus 3C (HRV3C) protease (Cordingley et al., 1990), a highly specific protease that recognizes the sequence LEVLFQGP, cleaving after Q and leaving a GP dipeptide at the start of the target. This allowed us to remove the target protein from the protein A-capture resin by protease treatment, avoiding low pH elution that is known to alter the conformations of type 1 fusion machine and could potentially alter the conformation of the probe. After the target sequence of interest, we incorporated a 10-residue linker followed by a biotin ligase-specific sequence (10lnQQ-AVI). This construct design enabled capture by protein A and on-column biotinylation (Figure 1B). As buffers for biotinylation and HRV3C cleavage are compatible, the entire process of biotinylation, HRV3C cleavage, and purification could in principle be accomplished with a single column, though we have generally added a polishing size exclusion chromatography step to ensure size homogeneity.

### Molecular Probe of Stabilized SARS-CoV-2 Spike Trimer with Biotinylation

To obtain a molecular probe of the SARS-CoV-2 spike, we incorporated stabilizing mutations into the spike target protein residues 14 to 1208 (Wrapp et al., 2020b), replacing RRAR at the S1/S2 cleavage site with GSAS and KV between domains HR1 and CH with PP (Figure 2 and Table S1) and adding the T4-phage fibritin trimerization domain (foldon) at the C-terminus (Efimov et al., 1994; Miroshnikov et al., 1998) (this construct is hereafter referred to as S2P), and expressed it by transient mammalian cell transfection. We observed cleavage by HRV3C protease to be impeded with the glutathione-S-transferase-tagged, but not the His-tagged version of the protease, suggesting partial steric constraints on access to the HRV3C site; overall we could obtain ∼0.5 mg of the purified S2P probe per liter cell culture after 24 h incubation at 4 °C with the His-tagged HRV3C.

**Figure 2.**
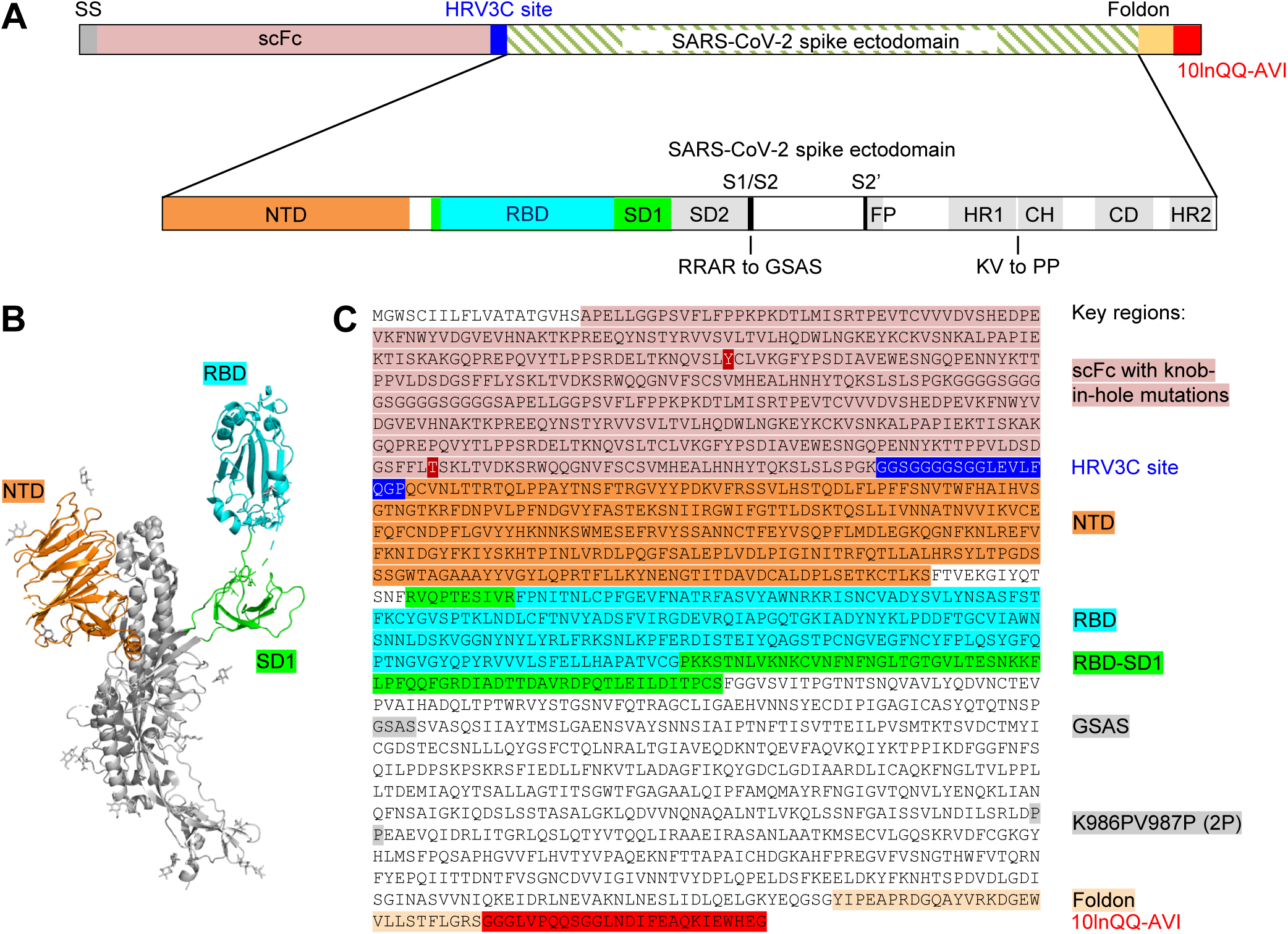
Design of the SARS-CoV-2 Spike S2P Molecular Probe. (A) Schematic of construction of the SARS-CoV-2 spike probe containing the ectodomain of the spike protein, the GSAS mutations replacing the RRAR furin cleavage site and the K986PV987P (2P) mutation stabilizing the spike in pre-fusion conformation, a foldon domain and a 10-amino acid linker (10lnQQ) followed by the AVI tag. (B) Structural model of a SARS-CoV-2 spike protomer with the NTD, RBD and SD1 domains highlighted. (C) Sequence of the SARS-CoV-2 spike probe with key regions annotated. See also Table S1.

The protease-liberated probe SARS-CoV-2 S2P ran primarily as a single peak on size-exclusion chromatography and a major band on SDS-PAGE (Figure 3A, B), appeared homogeneous by negative-stain EM (Figure 3C), and was recognized by the RBD-binding antibody CR3022 (Tian et al., 2020; Yuan et al., 2020). We also characterized binding to three antibodies from a SARS-1 convalescent donor S652: S652-109, which recognizes the RBD, S652-112, which recognizes an epitope in S2, and S652-118, which recognizes the NTD. All three of these antibodies bound well to the S2P probe, as did the ACE2 receptor, but not to soluble CD4 (D1D2-sCD4) or antibody VRC01 (Wu et al., 2010), which bound a control probe encompassing HIV-1 envelope glycoprotein trimer BG505 DS-SOSIP (Figure 3D). We note that the IgGs of CR3022, S652-109, S652-118 and S652-S112 showed stronger binding than corresponding antigen-binding fragment (Fab), potentially due to avidity. All of these properties suggested the SARS-CoV-2 S2P probe to be a good biological mimic of the prefusion SARS-CoV-2 spike ectodomain.

**Figure 3.**
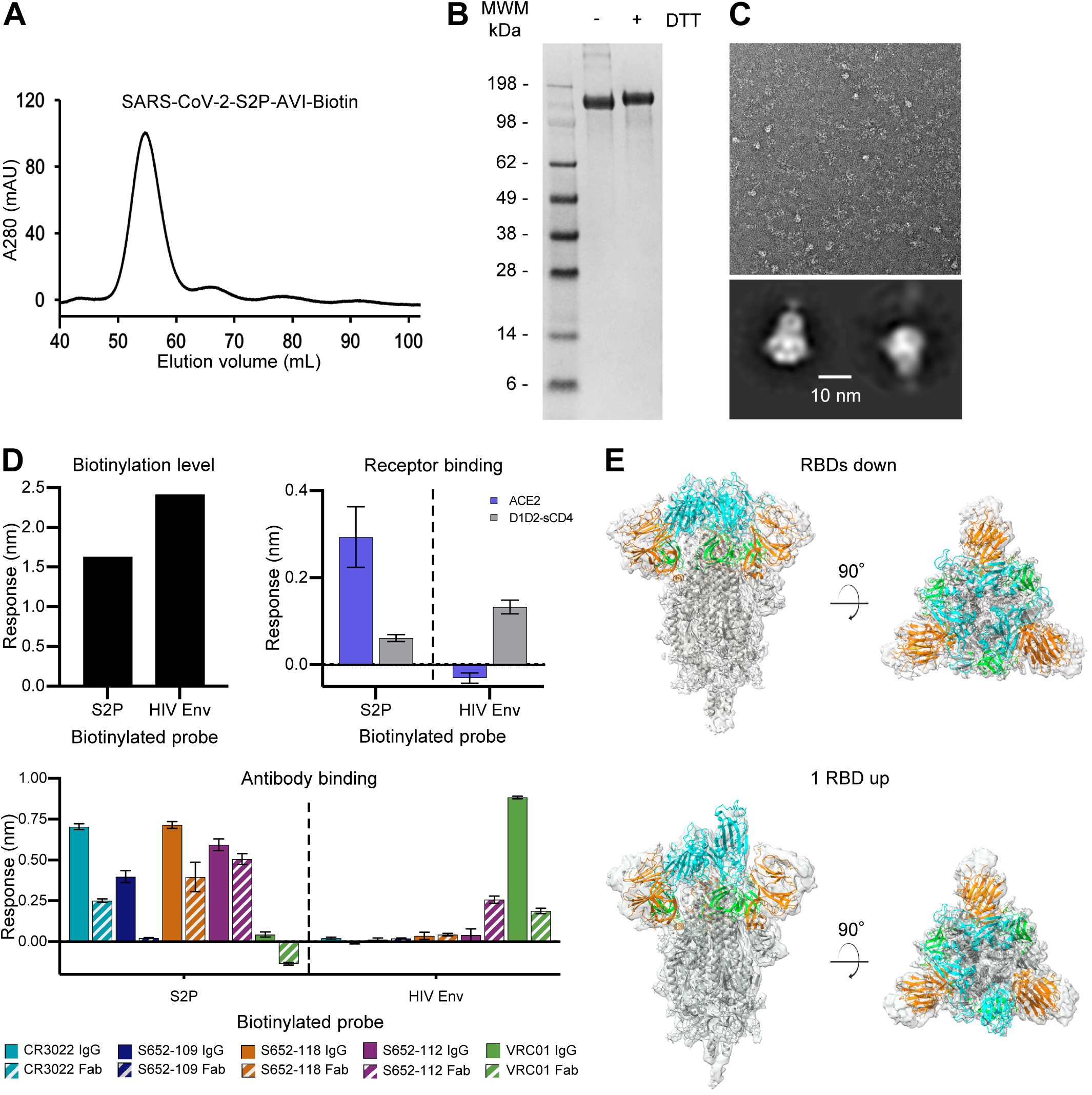
Physical Properties, Antigenic Characteristics, and Cryo-EM Structure of a Biotinylated SARS-CoV-2 S2P Probe. (A) Size exclusion chromatography of the biotinylated SARS-CoV-2 S2P probe in PBS buffer. (B) SDS-PAGE of the SARS-CoV-2 S2P probe with and without the reducing agent DTT. Molecular weight marker (MWM) was run alongside the probe. (C) Negative stain EM of the SARS-CoV-2 S2P probe. The 2D-class averages were shown below the wide-field raw EM image in 5-fold magnification. (D) Biotinylation, antigenicity and receptor recognition of the SARS-CoV-2 S2P probe. The level of biotinylation was evaluated by capture of probes at 40 μg/ml onto the streptavidin biosensors. A biotinylated HIV-1 Env stabilized in prefusion closed conformation by DS-SOSIP mutations and containing the same 10lnQQ-AVI tag at the C-terminus was used for comparison. Binding was assessed using 80 μg/ml S2P probe, 20 μg/ml HIV Env and receptors, or antibodies at 500 nM. Error bars represent standard deviation of triplicate measures. Low level non-specific binding of Fab S652-112 to the HIV-1 Env was observed. (E) Cryo-EM structures of biotinylated SARS-CoV-2 S2P probe. Domains are colored as in Figure 2C sequence. The C-terminal residues 1153-1208 plus the foldon, 10lnQQ-AVI tag, and biotin were not visible in the electron density. See also Figure S3 and Table S2.

### Cryo-EM Structure of a Biotinylated SARS-CoV-2 Spike Probe

To provide atomic-level insight into the mimicry of spike by probe, we determined the cryo-EM structure of the biotinylated SARS-CoV-2 S2P probe (Figures 3E and S3; Table S2). The cryo-EM grids were frozen with the biotinylated SARS-CoV-2 S2P probe in PBS at pH 7.4. We observed two distinct conformations, one with all three RBDs in the “down” position, displaying C3 symmetry, and the other with one RBD in the “up” position and the other two RBDs in the “down” position, C1-symmetric. We observed more particles (43,176) in the RBDs down conformation, which contributed to a 3.45 Å resolution map, whereas the 1 RBD up state (34,223 particles) 3D reconstruction achieved an overall resolution of 4.00 Å. The two final atomic models were very similar to published cryo-EM spike-ectodomain structures (Walls et al., 2020; Wrapp et al., 2020b). The C1-symmetric structure included residues 27-1147, while residues 27-1152 were modeled for the C3-symmetric structure; in both cases some regions could not be modeled because of their intrinsic flexibility. Five additional residues at the C-terminal region (1148-1152) were visible in the reconstruction density of the RBDs down structure, but - similar to published structures (Walls et al., 2020; Wrapp et al., 2020b) - the C-terminal region of the S2 domain (and in our case the rest of the probe) were not sufficiently ordered to be modeled. The two structures were similar except for the single RBD that was in the “up” position in the C1 symmetry structure. Excluding this single RBD, the RMSD was 0.95 Å between the two probe structures for 2738 aligned Cα atoms.

### Molecular Probes Comprising SARS-CoV-2 NTD, RBD and RBD-SD1 Regions

The domain structure of the SARS-CoV-2 spike was clearly evident in our cryo-EM structure and consistent with published accounts (Walls et al., 2020; Wrapp et al., 2020b). Based on these structures, we created constructs incorporating NTD, RBD and RBD-SD1 regions as separate molecular probes (Figure 4). Transient transfection yield for each of these constructs was greater than 5 mg/L. Both NTD and RBD eluted as a single peak on size-exclusion chromatography and exhibited a single band on SDS-PAGE (Figure 5A,B). RBD-SD1 eluted as two peaks on SEC, a disulfide-bonded dimer and a monomer (Figure 5A,B). We chose the monomeric form for further studies, as there is no biological basis for a higher association. Receptor recognition of these domain probes was specific for each construct, with ACE2 binding to RBD and RBD-SD1 probes, but not the NTD probe (Figure 5C, left). Similarly, antibodies CR3022 and S652-109 recognized RBD and RBD-SD1, while the S652-112 showed no binding to these two probes (Figure 5C, right); and antibody S652-118 recognized only the NTD probe. These results indicate that the truncated domain probes folded properly and retained the native conformation as on the full-length S2P probe.

**Figure 4.**
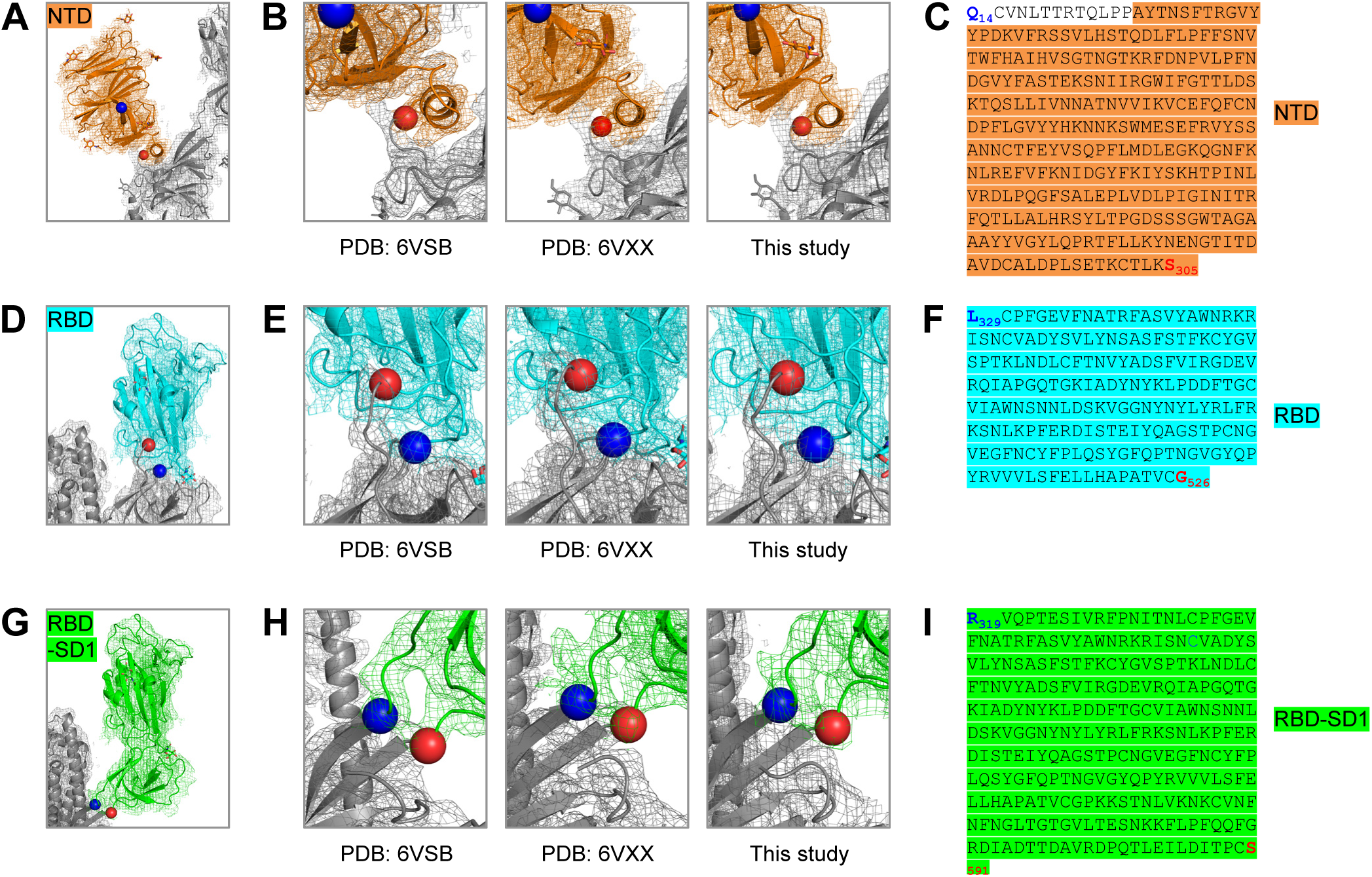
Structure-Based Definition of SARS-CoV-2 Molecular Probes Comprising the NTD, RBD and RBD-SD1 Domains. (A) Cryo-EM structure of the NTD domain in the S2P probe determined in this study (Figure 3E), with reconstruction density shown in orange for NTD domain, and gray otherwise. First ordered residue with density (A27) is highlighted with a blue sphere; last residue of NTD domain (S305) is highlighted with a red sphere. (B) Close-up view of the NTD termini. (C) Sequence of NTD domain probe. The sequence is highlighted in orange except for residues 14-26, which are disordered in the cryo-EM structures. (D) Cryo-EM structure of the RBD domain in spike (Figure 3E), with reconstruction density shown in cyan for RBD domain, and gray otherwise. First residue with density (L329) is highlighted with a blue sphere; last ordered residue of RBD domain (G526) is highlighted with a red sphere. (E) Close-up view of the spike RBD termini. (F) Sequence of RBD domain probe highlighted in cyan. (G) Cryo-EM structure of the RBD-SD1 domains in spike (Figure 3E), with reconstruction density shown in green for RBD-SD1 domain, and gray otherwise. First residue with density (R319) is highlighted with a blue sphere; last ordered residue of RBD-SD1 domain (S591) is highlighted with a red sphere. (H) Close-up view of the spike RBD-SD1 termini. (I) Sequence of the RBD-SD1 domain probe highlighted in green. See also Figures S1 and S3 and Tables S1 and S2.

**Figure 5.**
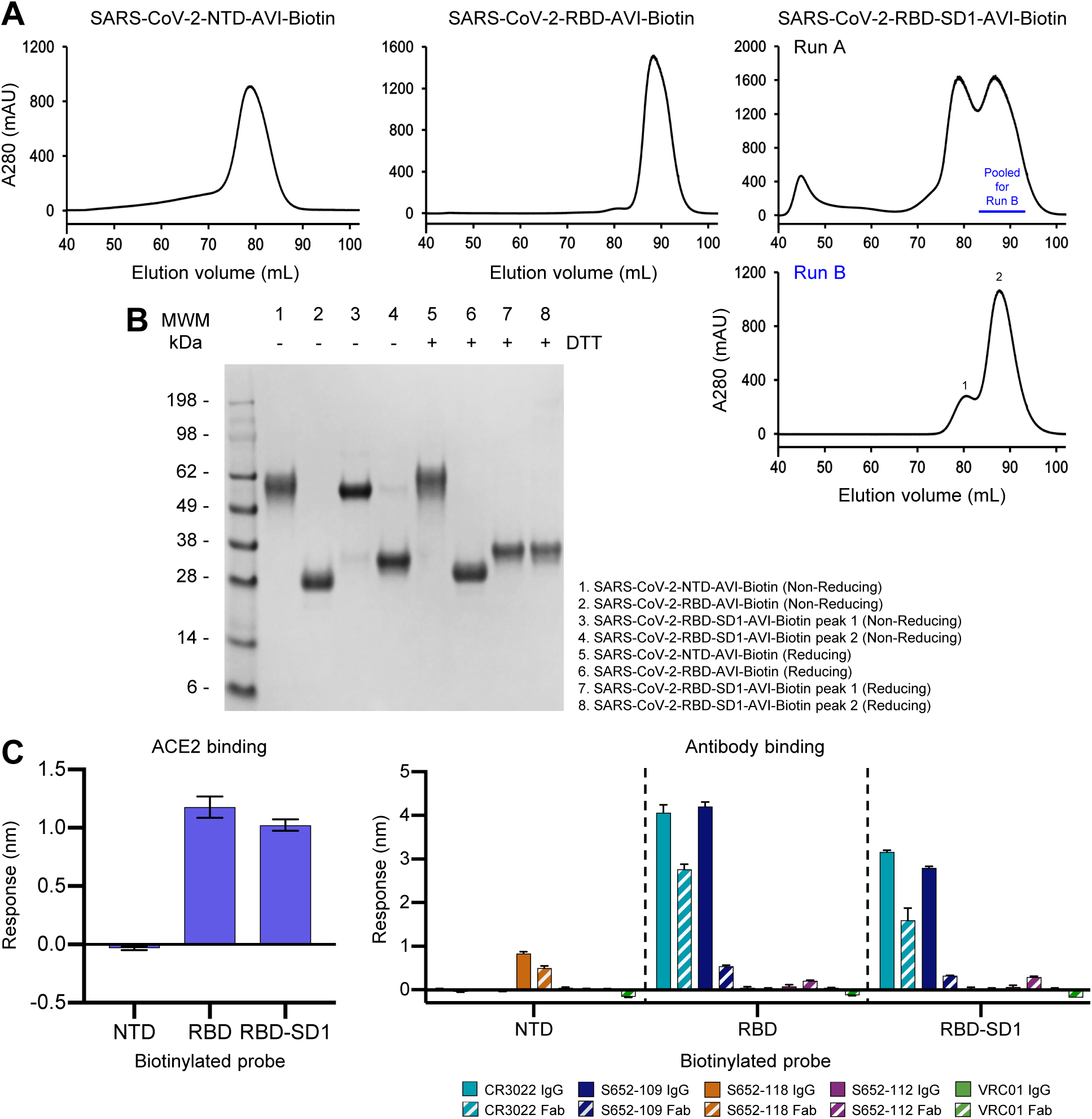
Physical Properties and Antigenic Characteristics of Molecular Probes Comprising SARS-CoV-2 NTD, RBD, and RBD-SD1 Domains. (A) Size exclusion chromatography of biotinylated NTD, RBD and RBD-SD1 molecular probes. (B) SDS-PAGE of NTD, RBD and RBD-SD1 molecular probes. RBD-SD1 peak 1 appeared to be partially disulfide linked, and was therefore removed from further analysis; RBD-SD1 thus refers to the monomeric peak 2. (C) Receptor binding (left) and antibody binding (right). Both RBD and RBD-SD1 probes bound to the ACE2 receptor and antibody S652-109 while the NTD probe interacted with antibody S652-118 only. Binding was assessed using 5 μg/ml probe and 500 nM receptor or antibody. Error bars represent standard deviation of triplicate measures. Low level non-specific binding of S652-112 Fab to RBD and RBD-SD1 was observed. See also Figure S2.

### Molecular Probes Comprising Specific Knockouts of ACE2 Interaction with RBD

Structural information can define not only domain boundaries, but also biological interfaces. Based on the structure of the RBD domain with ACE2 (Lan et al., 2020) (Figure 6A), we designed constructs with ACE2-interface residues mutated to Arg to knockout ACE2 affinity (Figure 6B). We expressed and purified two double-Arg mutants (RBD-L455RA475R-AVI and RBD-L455RG496R-AVI) and one triple-Arg mutant (RBD-L455RA475RG502R-AVI). All three eluted in a single peak on size-exclusion chromatography and exhibited a single band on SDS-PAGE (Figure 6C,D). All three mutants completely ablated ACE2 binding as designed (Figure 6E, left). By contrast, CR3022 recognition by all three mutants was mostly retained as the epitope of CR3022 and ACE2-binding site are located at different locations on RBD; S652-109 IgG recognition by the double-Arg mutants was mostly retained and its IgG recognition by RBD-L455RA475RG502R-AVI was reduced but at detectable level, indicating a potential epitope overlapping the G502 region. Similar to observations with S2P-AVI, RBD-AVI and RBD-SD1-AVI, S652-109 Fab recognition by all three mutants was substantially reduced (Figure 6E, right). Our results indicated that the ACE2-interface-mutant probes can be used to specifically distinguish antibodies targeting the ACE2-binding site and other epitopes on the RBD.

**Figure 6.**
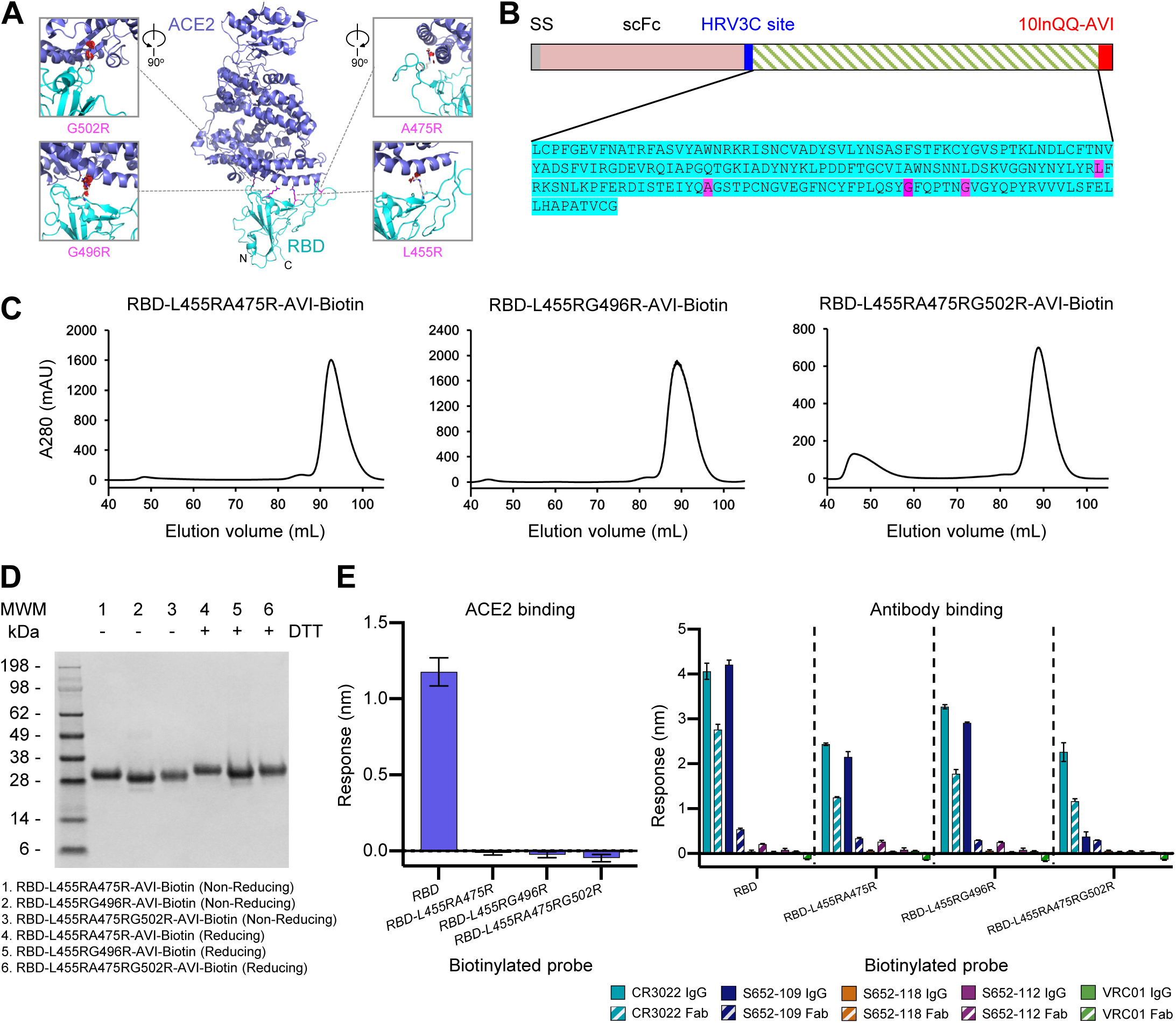
Molecular Probes Comprising Specific Mutations for Knocking out ACE2 Interaction with RBD. (A) Structural model of the RBD-ACE2 complex, with inset panels highlighting RBD mutations designed to reduce ACE2 recognition. ACE2 and RBD are shown in cartoon representation and colored blue and cyan, respectively. Arg mutations are shown in stick representation, with potential clashes with ACE2 highlighted with red discs. (B) Schematic and sequence of mutant RBD probes with sites mutated to Arg highlighted in magenta. (C) Size exclusion chromatography of RBDs with mutations altering ACE2 binding. (D) SDS-PAGE of mutant RBD probes with and without reducing agent as indicated. (E) Receptor and antibody binding of mutant RBD probes. All the knockout mutations abolished RBD binding to the ACE2 receptor while retaining binding to both CR3022 and S652-109 antibodies. Binding was assessed using 7.5 μg/ml RBD-L455RA475R, 80μg/ml RBD-L455RG496R, or 15μg/ml (A) (A) (E) (E) (F) RBD-L455RA475RG502R to account for varying biotinylation levels among the mutants. Receptor or antibodies were at 500 nM. Error bars represent standard deviation of triplicate measures. See also Figure S2 and Table S1.

### Validation of Biotinylated Probes

To assess the molecular probes for their quality and function, we first performed kinetic analyses of ACE2 receptor and antibodies binding to the probes by SPR (Figure 7A). Purified biotinylated probes were immobilized on streptavidin sensor chip surface, and ACE2 and antibodies were in the mobile phase. ACE2 bound the full-length ectodomain probe S2P with a K_D_ of 46.5 nM, and its affinity to the isolated RBD domain probe was somewhat lower with a K_D_ of 121 nM. We found the S2P probe to be sensitive to chip regeneration required to remove antibody, so we used single-cycle kinetics to analyze the interaction with the Fabs of antibodies CR3022, S652-109, S652-112 and S652-118. These all bound SARS-CoV-2 S2P probe with high affinity (Figure 7A, top row). Interestingly, the binding on-rates for all three of the antibodies were comparable, but the off-rates varied. CR3022, an antibody known to bind RBD but not neutralize the SARS-CoV-2 virus (Yuan et al., 2020), bound to SARS-CoV-2 S2P at a lower affinity than ACE2 (91.4 vs 46.5 nM, respectively), although CR3022 had a higher affinity to the RBD probe (25.0 nM) than to S2P (91.4 nM). Another RBD-binding non-neutralizing antibody, S652-109, bound to the RBD and S2P probes at 1.63 and 3.96 nM, respectively. Antibody S652-112, also non-neutralizing, did not bind NTD or RBD probes (Figure 5C), but did bind S2P probe with a K_D_ of 0.32 nM, consistent with it being S2 directed. An NTD-specific antibody, S652-118, which does have some neutralizing capacity, bound NTD and S2P probes at and 1.46 nM, respectively. These results confirm that the biotinylated probes have the binding specificity for the receptor and antibodies and are functioning properly as expected.

**Figure 7.**
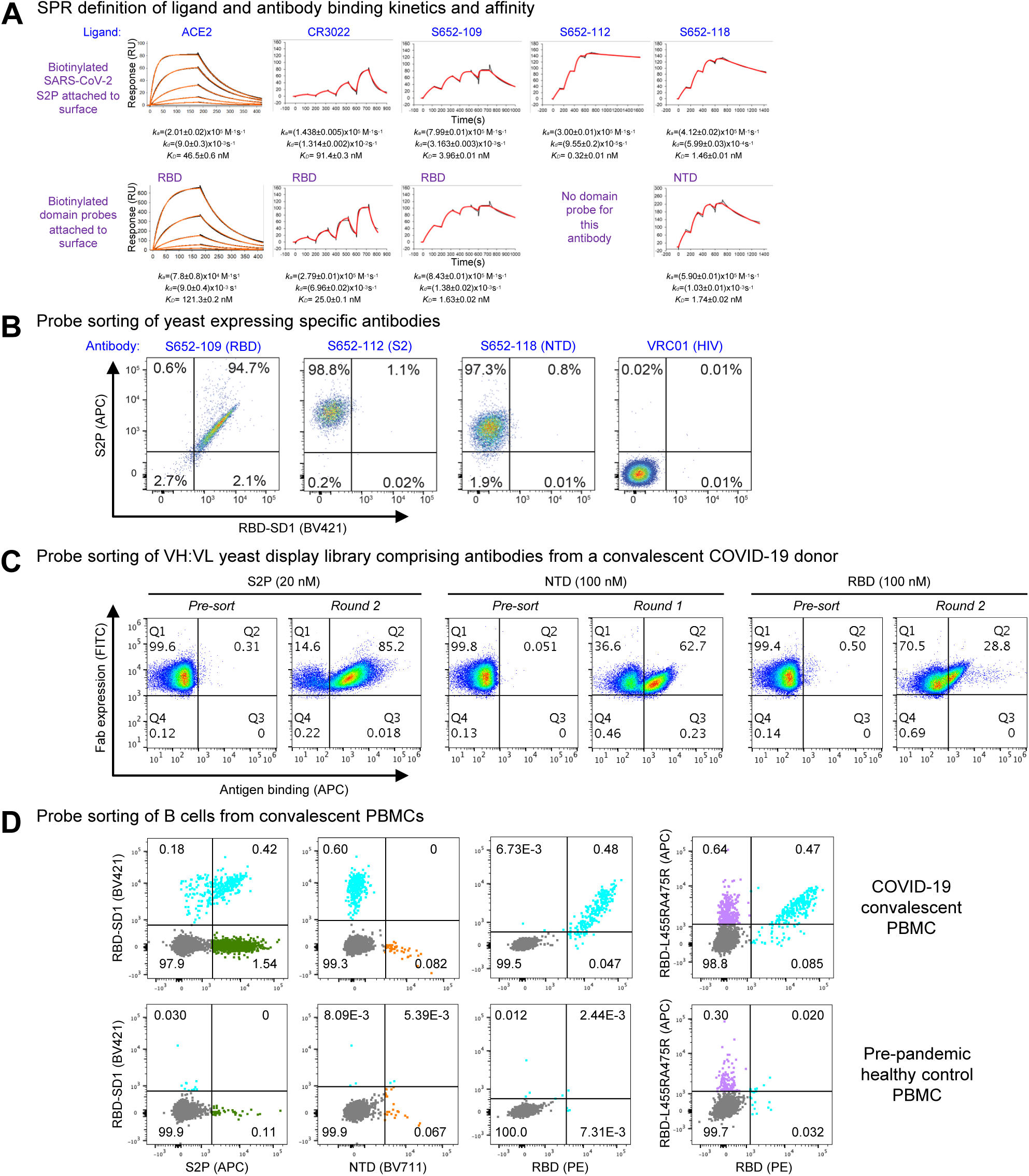
Validation of Biotinylated Probes. (A) SPR standard kinetic experiments for ACE2 and single-cycle kinetic experiments for Fabs CR3022, S652-109, S652-112 and S652-118 binding over biotinylated spike (top row) and biotinylated-RBD or -NTD (bottom row) loaded onto a streptavidin sensorchip. Black traces represent the experimental data and the red traces represent the fit to a 1:1 interaction model. The error in each measurement represents the error of the fit. (B) Binding of yeast expressing SARS-CoV cross-reactive Fabs to SARS-CoV-2 antigenic probes. HIV envelope-targeting VRC01 Fab was used as a negative control. (C) Sorting of yeast cells encoding antibody transcripts from a convalescent COVID-19 donor with S2P, NTD and RBD-based probes. (D) Sorting of B cells from COVID-19 convalescent PBMCs. (Top row) PBMCs obtained from a SARS-CoV-2-infected subject 75 days after symptom onset were stained with surface antibodies and probes as outlined in the methods to allow for identification and specificity-determination of SARS-CoV-2 spike-specific IgG+ and IgA+ B cells. (Bottom row) PBMCs obtained pre-pandemic from a healthy donor serve as a control for background binding of the probes. See also Figures S4-S7.

We next tested the probes for sorting antigen-specific yeast cells displaying SARS-CoV-2 antibodies (Figures 7B, S4 and S5). We expressed antibodies S652-109, S652-112, and S652-118 on yeast cell surface. The biotinylated probes, S2P, RBD-SD1 and the ACE2-knockout triple-Arg mutant RBD-L455RA475RG502R, were freshly labeled with fluorochromes and used to sort the yeast cells. As expected, the S2P probe recognized all cells expressing S652-109, S652-112 and S652-118, but not those with the control antibody, whereas the RBD-SD1 probes recognized only cells expressing S652-109 (Figure 7B and S5). The triple-Arg mutant RBD-L455RA475RG502R also recognized S652-109 (Figure S5), consistent with the BLI binding assay above that showed residual binding was retained (Figure 6E). These results confirm the utility of the probes for sorting cells expressing SARS-CoV-2 spike specific antibodies on cell surface.

Having established the cell-sorting utility of the probes, we next set out to test the probes in sorting and enriching natively paired VH:VL yeast display libraries of convalescent COVID-19 donors for SARS-CoV-2 spike-specific antibodies (Figures 7C and S6) (Wang et al., 2018a). Depending on the initial level of antigen-specific antibodies present, just one or two rounds of sorting and enrichment could increase the antigen-specific cells by several orders of magnitude, indicating substantial utility of the probes for cell sorting.

To further demonstrate the utility of the probes in sorting of human B cells, we obtained peripheral blood mononuclear cells (PBMCs) from a convalescent SARS-CoV-2 patient and sorted the cells with fluorochrome-labeled probes (Figures 7D and S7). S2P-positive, RBD-SD1-positive, and RBD-positive B cells were identified from the convalescent patient PBMCs but not from pre-pandemic healthy control PBMCs (Figure 7D, left three panels). The double-Arg mutant RBD-L455RA475R, by contrast, recognized only about 2-fold more B cells than the control (Figure 7D, far right panel), with only a small proportion of B cells not binding mutant versus RBD-SD1 (Figure 7D, comparing 3^rd^ and 4^th^ panels).

Overall, the designed probes were found to be functional in SPR and in sorting yeast cells or B cells with spike-reactive antibodies.

## Discussion

Molecular probes comprising key targets such as the SARS-CoV-2 spike and its domains can be of broad utility. For vaccine development, such probes can be used to define and monitor the elicited responses, and for antibody characterization, such probes can serve as critical molecular tags, facilitating the identification of the highly effective antibodies which could be used for passive therapy (Cao et al., 2020; reviewed in Casadevall et al., 2004; Graham and Ambrosino, 2015; Walker and Burton, 2018). In diagnostics, they can be employed to assess sera reactivity and to provide sensitive markers of infection (Perera et al., 2020; Yan et al., 2020b; Zhao et al., 2020). In pathogenesis, they can assist in delineating susceptible cells that virus might infect or in tracking viral variants, including those that might have selective advantages (Korber et al., 2020). Here we show how structure-based design coupled to incorporation of (i) a purification-tag based construct, (ii) a sequence-specific protease, and (iii) a sequence-specific biotin ligase, enable streamlined probe development.

For probe construction, we used a process incorporating the ‘cut-and-paste’ assembly of necessary components, employing an N-terminal purification tag and sequence stretches targeted by sequence-specific enzymes: the HRV3C protease and biotin ligase. Other sequence-specific cleavage sites such as thrombin could be incorporated, to further control the presence of components surrounding the target protein of interest. Such protease-specific cleavage, however, can lead to the addition of residual amino acids at resultant N- and C-termini after protease cleavage; we note in this context that an N-terminal HRV3C site appends only a dipeptide, Gly-Pro, to the N-terminus of the liberated target. Importantly, structure of the target protein can define domain boundaries, allowing for separate probes of specific domains or domain combinations.

The structure-based methods we describe here for probe construction may allow for the assessment of immune responses of other type 1 fusion machines. We previously created hemagglutinin trimer probes with a Y96F mutation (Whittle et al., 2014) and used them to identify broadly neutralizing antibodies against influenza (Joyce et al., 2016). We also created Ebolavirus trimer probes that could be used to identify protective neutralizing antibodies against Sudan outbreak Ebolavirus strain (Corti et al., 2016; Gaudinski et al., 2019; Misasi et al., 2016). And we previously used a biotinylated envelope ‘SOSIP’ probe to identify broadly neutralizing, fusion peptide-directed antibodies against HIV-1 (Kong et al. Science 2016). Overall, the cut-and-paste structure-based design described here should be easily adapted to the streamlined development of molecular probes against not only these pathogens, but also emerging pathogens, as shown here for SARS-CoV-2.

## Acknowledgments

We thank R. Grassucci, Y.-C. Chi and Z. Zhang from the Cryo-EM Center at Columbia University for assistance with cryo-EM data collection, M. G. Joyce for antibody CR3022, J. Stuckey for assistance with figures, and members of the Virology Laboratory and Vector Core, Vaccine Research Center, for discussions and comments on the manuscript. Support for this work was provided by the Intramural Research Program of the Vaccine Research Center, National Institute of Allergy and Infectious Diseases (NIAID). Support for this work was also provided by COVID-19 Fast Grants, the Jack Ma Foundation, the Self Graduate Fellowship Program, and NIH grants DP5OD023118, R21AI143407, and R21AI144408. Some of this work was performed at the Columbia University Cryo-EM Center at the Zuckerman Institute, and some at the Simons Electron Microscopy Center (SEMC) and National Center for Cryo-EM Access and Training (NCCAT) located at the New York Structural Biology Center, supported by grants from the Simons Foundation (SF349247), NYSTAR, and the NIH National Institute of General Medical Sciences (GM103310).

## Author Contributions

TZ designed the probes; TZ, ITT and ASO produced the probes; GC and JG determined cryo-EM structure of S2P; TZ and AN performed Octet antigenic assessment; WS provided ACE2; YT performed negative-stain EM; LW, KSC, and YZ provided antibodies S652-109, -112 and -118; SW assisted with data analysis and manuscript assembly; BZ produced antibody CR3022; PSK carried out SPR; YP, JM and NJS performed monoclonal yeast display analysis; BBB, ASF, LL, SNLA, BM, MOdS, XP, PW, JRW, MY, DDH and BJD carried out natively paired antibody yeast display analysis; EP, AD, LC, OA, KSC, TR and BSG preformed PBMC B cell staining analysis; JRM supervised ACE2 and antibodies S652-109, -112 and -118 production; LS supervised SPR and cryo-EM structural analyses; TZ, LS and PDK oversaw the project and – with ITT, ASO, GC, JG, YT, SW, JM, BJD, and TR – provided a first draft of the manuscript, with all authors providing revisions and comments.

## STAR□METHODS

- KEY RESOURCES TABLE
- CONTACT FOR REAGENT AND RESOURCE SHARING
- EXPERIMENTAL MODEL AND SUBJECT DETAILS
  - Patient samples
  - Cell lines
- METHOD DETAILS
  - Expression plasmid construction for SARS-CoV-2 probes
  - SARS-CoV-2 probe preparation
  - Expression and preparation of the ACE2 receptor
  - Expression and preparation of antibodies
  - Bio-Layer Interferometry
  - Surface plasmon resonance
  - Negative-stain electron microscopy
  - Cryo-EM data collection and processing
  - Cryo-EM model fitting
  - Probe conjugation
  - Monoclonal yeast display analysis
  - Experimental sorting of natively paired antibody heavy:light yeast display libraries
  - PBMC B cell stain
- QUANTIFICATION AND STATISTICAL ANALYSIS
- DATA AND CODE AVAILABILITY

### KEY RESOURCES TABLE

| REAGENT or RESOURCE | SOURCE | IDENTIFIER |
| --- | --- | --- |
| **Antibodies** |  |  |
| CR3022 | Yuan et al., 2020 | N/A |
| S652-109 | This study | N/A |
| S652-112 | This study | N/A |
| S652-118 | This Study | N/A |
| VRC01 | Wu et al., 2010 | N/A |
| **Bacterial and Virus Strains** |  |  |
| None |  |  |
| **Chemicals, Peptides, and Recombinant Proteins** |  |  |
| Superdex200 10/300GL Column | GE Healthcare Life Sciences | Cat# 28990944 |
| MabSelect SuRe Protein A Resin | GE Healthcare Life Sciences | Cat# 17543802 |
| SARS-CoV-2-S2P-AVI | This study | N/A |
| SARS-CoV-2-NTD-AVI | This study | N/A |
| SARS-CoV-2-RBD-AVI | This study | N/A |
| SARS-CoV-2-RBD-SD1-AVI | This study | N/A |
| SARS-CoV-2-RBD-L455RA475R-AVI | This study | N/A |
| SARS-CoV-2-RBD-L455RG496R-AVI | This study | N/A |
| SARS-CoV-2-RBD-L455RA475RG502R-AVI | This study | N/A |

| **Critical Commercial Assays** |  |  |
| --- | --- | --- |
| Turbo293™ Transfection Kit | ThermoFisher Scientific Inc. | Cat# A14525 |
| BirA biotin-protein ligase bulk reaction kit | Avidity | BirA500 |
| **Deposited Data** |  |  |
| Cryo-EM map: SARS-CoV-2 S2P C1 symmetry | EMDB | EMDB: EMD-22162 |
| Cryo-EM structure: SARS-CoV-2 S2P C1 symmetry | PDB | PDB: 6XF6 |
| Cryo-EM map: SARS-CoV-2 S2P C3 symmetry | EMDB | EMDB: EMD-22161 |
| Cryo-EM structure: SARS-CoV-2 S2P C3 symmetry | PDB | PDB: 6XF5 |
| **Experimental Models: Cell Lines** |  |  |
| Expi293F cells | ThermoFisher Scientific Inc | Cat# A14527 |
| FreeStyle 293-F cells | ThermoFisher Scientific Inc | Cat# R79007 |
| **Recombinant DNA** |  |  |
| pVRC8400 vector | https://www.addgene.org | Cat# 63160 |
| pVRC8400-CR3022 | Yuan et al., 2020 | N/A |
| pVRC8400-S652-109 | This study | N/A |
| pVRC8400-S652-112 | This study | N/A |
| pVRC8400-S652-118 | This study | N/A |
| pVRC8400-SARS-CoV-2-S2P-AVI | This study | N/A |
| pVRC8400-SARS-CoV-2-NTD-AVI | This study | N/A |
| pVRC8400-SARS-CoV-2-RBD-AVI | This study | N/A |
| pVRC8400-SARS-CoV-2-RBD-SD1-AVI | This study | N/A |
| pVRC8400-SARS-CoV-2-RBD-L455RA475R-AVI | This study | N/A |
| pVRC8400-SARS-CoV-2-RBD-L455RG496R-AVI | This study | N/A |
| pVRC8400-SARS-CoV-2-RBD-L455RA475RG502R-AVI | This study | N/A |
| **Software and Algorithms** |  |  |
| GraphPad Prism Software | GraphPad Prism Software, Inc. | N/A |
| Scrubber 2.0 | BioLogic Software | http://www.biologic.com.au/scrubber.html |
| The PyMol Molecular Graphics System, v1.8.6 | Schrödinger, LLC | https://pymol.org/2/ |
| BLI Acquisition & Analysis Software | ForteBio | https://www.fortebio.com/products/octet-systems-software |
| Leginon | Suloway et al., 2005 | https://sbgrid.org/software/titles/leginon |
| The PEP-Fold server | Shen et al., 2014<br>Thévenet et al., 2012 | https://bioserv.rpbs.univ-paris-diderot.fr/services/PEP-FOLD/ |
| GCTF | Zhang, 2016 | https://www.mrc-lmb.cam.ac.uk/kzhang/Gctf/ |
| CTFFind4 | Grigorieff, 2015 | http://grigoriefflab.janelia.org/ctffind4 |
| Dog Picker | Voss et al., 2009 | https://omictools.com/dog-picker-tool |
| RELION | Scheres, 2012 | https://www3.mrc-lmb.cam.ac.uk/relion/index.php/Main_Page |
| cryoSPARC | Punjani et al., 2017 | https://cryosparc.com |
| EMRinger | Barad et al., 2015 | http://emringer.com/ |
| Phenix | Adams et al., 2004 | https://sbgrid.org/software/ |
| Coot | Emsley and Cowtan, 2004 | https://sbgrid.org/software/ |
| UCSF Chimera | Pettersen et al., 2004 | https://www.cgl.ucsf.edu/chimera/ |

### RESOURCE AVAILABILITY

#### Lead Contact

Further information and requests for resources and reagents should be directed to and will be fulfilled by Peter D. Kwong.

#### Materials Availability

Plasmids for the SARS-CoV-2 probes with AVI tag generated in this study are available upon request for non-commercial research purposes.

#### Data and Code Availability

Cryo-EM maps of the biotinylated SARS-CoV-2 S2P have been deposited to the EMDB with accession codes EMD-22161 (RBDs down) and EMD-22162(1 RBD up), and fitted coordinates have been deposited to the PDB with accession codes 6XF5 and 6XF6, respectively.

### EXPERIMENTAL MODEL AND SUBJECT DETAILS

#### Patient samples

Peripheral blood mononuclear cells (PBMCs) for B cell sorting were obtained from a convalescent SARS-CoV-2 patient (collected 75 days post symptom onset under an IRB approved clinical trial protocol, VRC 200 – ClinicalTrials.gov Identifier: NCT00067054) and a healthy control donor from the NIH blood bank pre-SARS-CoV-2 pandemic.

PBMCs for experimental sorting of natively paired antibody heavy:light yeast display libraries were obtained from 2 participants enrolled in the COVID-19 Cohort Study at Columbia University Irving Medical Center. Participants were both men in their 50’s, hospitalized for severe respiratory decompensation requiring mechanical ventilation after confirmed SARS-CoV-2 infection. Specimens were obtained at 25 days (1605) and 32 days (NYP01) after symptom onset. Patient samples were collected after obtaining informed consent and appropriate Institutional Review Board approval (IRB#AAAS9517).

#### Cell lines

Expi293F and FreeStyle 293-F cells were purchased from Thermo Fisher Scientific. The cells were used directly from the commercial sources following manufacturer suggestions as described in detail below.

### METHOD DETAILS

#### Construction of expression plasmids for SARS-CoV-2 probes

DNA sequences coding for the SARS-CoV-2 spike or its domains were cloned into a pVRC8400-based expression vector by GeneImmune. The gene of interest was placed after the DNA sequence encoding a N-terminus single chain human Fc with knob-in-hole mutations, a “GGSGGGGSGG” linker and an HRV3C protease cleavage site for purification with protein A column and on-column cleavage. DNA sequence encoding a C-terminus “GGGLVPQQSG” linker named10lnQQ and an AVI tag was placed after the gene of interest for biotinylation (Table S1).

For the scFc-SARS-CoV-2 S2P-AVI construct, the cloned insertion included the spike protein residues 14 to 1208 with GSAS replacing the RRAR at the S1/S2 cleavage site and K986P and V987P stabilization mutations, a GSG linker and the T4-phage fibritin trimerization domain (foldon) as described by Wrapp and colleagues (Wrapp et al., 2020b). For the scFc-SARS-CoV-2 NTD-AVI construct, spike gene encoding residues 14-305 was cloned. For the scFc-SARS-CoV-2 RBD-AVI construct, spike gene encoding residues 329-526 was cloned. For the scFc-SARS-CoV-2 RBD-SD1-AVI construct, spike gene encoding residues 319-591 was cloned. The constructs comprising specific mutations for knocking out ACE2 interaction with RBD were made by site directed mutagenesis in the scFc=SARS-CoV-2-RBD-AVI plasmid, either in double Arg substitutions at L455 and A475 (L455RA475R), or at L455 and G496 (L455RG496R), or in triple Arg substitutions at L455, A475, and G502 (L455RA475RG502R). A model of the 10lnQQ linker and AVI tag was obtained on the PEP-FOLD server (Shen et al., 2014; Thevenet et al., 2012) to show the location of this C-terminal addition in Figure S1.

#### Expression and preparation of SARS-CoV-2 probes

Single chain Fc tagged SARS-CoV-2 probes were produced by transient transfection of 293 Freestyle cells as previously described (Wrapp et al., 2020b). One liter of cells was transfected with 1 mg of SARS-CoV-2 probe plasmid, pre-mixed with 3 ml of Turbo293 transfection reagent. The cells were allowed to grow for 6 days at 120 rpm, 37 °C, and 9 % CO_2_, after which the supernatant was harvested by centrifugation and filtered. The cleared supernatant was incubated with 5ml of PBS-equilibrated Protein A resin for two hours, after which the resin was then collected and washed with PBS. The captured SARS-CoV-2 probe was then concurrently biotinylated with 10-80 μg BirA enzyme (∼2.5 μg per 10 nmol AVI substrate) and 100 μM biotin, as well as cleaved from the Fc purification tag with 200 μg of HRV3C, which was prepared as described (Antoniou et al., 2017), in a reaction mixture containing 10 mM Tris pH 8.0, 50 μM bicine, 10 mM ATP, 10 mM Mg(OAc)_2_. For RBD and NTD domains, addition of 100 μM DTT in the reaction may facilitate the cleavage but it is optional. After incubation at 4°C overnight, the cleaved protein was collected, concentrated and applied to a Superdex 200 16/60 gel filtration column equilibrated with PBS. Peak fractions were pooled and concentrated to 1 mg/ml.

#### Expression and preparation of the ACE2 receptor

The human ACE2 (1-620aa) expression plasmid was constructed by placing an HRV3C cleavage site, a monomeric Fc tag and an 8xHisTag at the 3’-end of the ACE2 gene. After transient transfection of 293F cells, the cell culture supernatant was harvested 4 days post transfection and loaded onto a Protein A affinity column. The Fc-tagged protein was eluted with IgG elution buffer and then dialyzed against PBS before overnight HRV3C digestion at 4 °C. Cleaved protein was passed over a protein A column to remove the Fc tag. ACE2 from the protein A column flow through was further purified with a Superdex 200 16/60 column in 5 mM HEPES, pH7.5 and 150 mM NaCl.

#### Expression and preparation of antibodies

DNA sequences of antibody heavy and light chain variable regions of antibody CR3022 (Yuan et al., 2020) and of donor S652 antibodies, S652-109, S652-112, and S652-118, were synthesized and subcloned into the pVRC8400 vector, as described previously (Wu et al., 2011). Antibody expression was carried out by co-transfection of both heavy and light chain plasmids in Expi293F cells (Thermo Fisher) using Turbo293 transfection reagent (Speed BioSystems). On day 6 post transfection, the culture supernatant was harvested and loaded on a protein A column. After washing with PBS for 3 column volumes, IgG was eluted with an IgG elution buffer (Pierce) and immediately neutralized by addition of one tenth volume of 1M Tris-HCl pH 8.0. The Fab fragments were generated by overnight Endoproteinase LysC (New England Biolabs) digestion at 37°C with a LysC:IgG ratio of 1:2000 (w/w) and passed through a protein A column to remove uncut IgG and Fc fragments.

#### Bio-Layer Interferometry

A FortéBio Octet HTX instrument (FortéBio) was used to analyze biotinylation level, antigenicity, and receptor recognition of the molecular probes. Assays were performed at 30°C in tilted black 384-well plates (Geiger Bio-One) in PBS + 0.02% Tween20, 0.1% BSA, 0.05% sodium azide with agitation set to 1,000 rpm. Biotinylated SARS-CoV-2 spike (80 μg/ml), NTD (40 μg/ml), RBD (5 μg/ml), RBD-SD1 (5 μg/ml), RBD-L455RA475R (5 μg/ml), RBD-L455RG496R (80 μg/ml), RBD-L455RA475RG502R (15 μg/ml) and HIV Env (20 μg/ml) were loaded on to streptavidin biosensors for 150 seconds. Binding to biotinylated probes was measured by dipping immobilized probes into solutions of receptors and antibodies at 500 nM for 180 seconds. Receptor and antibody binding were assessed in triplicate. Parallel correction to subtract systematic baseline drift was carried out by subtracting the measurements recorded for a loaded sensor dipped into buffer only control well. The reference corrected response values are reported as averages of maximum signal during the association step.

#### Surface plasmon resonance

SPR binding experiments were performed using a Biacore T200 biosensor, equipped with either a Series S SA chip for the Fab binding assays, or a Series S CM4 chip immobilized with NeutrAvidin, for the ACE2 binding assays. All measurements were performed at 25°C in a running buffer of 10 mM HEPES pH 7.4, 150 mM NaCl, and 0.05% (v/v) Tween-20.

Biotinylated spike, RBD, and NTD were captured over independent flow cells at 1000-1100 RU, 300-500 RU and 260 RU, respectively. ACE2-analyte samples were prepared in running buffer at a concentration range of 300-1.2nM using a three-fold dilution series. Binding over the spike and RBD or NTD surfaces as well as over a neutravidin reference surface was tested for 180s followed by a 300s dissociation phase at 50μL/min. Blank buffer cycles were performed throughout the experiment by injecting running buffer instead of ACE2 to subtract systematic noise from the binding signal. The data was processed and fit to 1:1 binding model using Scrubber 2.0 (BioLogic Software).

To avoid the need for surface regeneration that arises with the low off-rate Fabs, we used single-cycle kinetics for the Fab binding experiments. Each Fab was tested at analyte concentrations 180-6.66nM prepared in running buffer using a three-fold dilution series. Binding over the spike or RBD surface or the NTD surface for S652-118, was monitored for 120s, followed by a dissociation phase of 120s-900s depending on the interaction. Four blank buffer single cycles were performed by injecting running buffer instead of Fab to remove systematic noise from the binding signal. The data was processed and fit to 1:1 single cycle model using the Scrubber 2.0 (BioLogic Software).

#### Negative-stain electron microscopy

Purified SARS-CoV-2 S samples were diluted to approximately 0.02 mg/ml with a buffer containing 10 mM HEPES, pH 7, and 150 mM NaCl. A 4.7-µl drop of the diluted sample was applied to a glow-discharged carbon-coated copper grid. The grid was washed with the same buffer, and adsorbed protein molecules were negatively stained with 0.75% uranyl formate. Micrographs were collected at a nominal magnification of 100,000x (pixel size: 0.22 nm) using SerialEM (Mastronarde, 2005) on an FEI Tecnai T20 electron microscope operated at 200 kV and equipped with an Eagle CCD camera. Particles were picked automatically using in-house written software (Y.T., unpublished). Reference-free 2D classification was performed with Relion 1.4 (Scheres, 2012).

#### Cryo-EM data collection and processing

Biotinylated SARS-CoV-2 spike probe was diluted to a final trimer concentration of 0.33 mg/mL in PBS, pH 7.4. The sample (2 µL) was applied to a glow-discharged carbon-coated copper grid (CF 1.2/1.3) and vitrified using a Vitrobot Mark IV with a wait time of 30 seconds and a blot time of 3 seconds.

Data were collected using the Leginon software (Suloway et al., 2005) installed on a Titan Krios electron microscope operating at 300 kV, equipped with a Gatan K3-BioQuantum direct detection device. The total dose was fractionated for 3 s over 60 raw frames. Motion correction, CTF estimation, particle picking and extraction, 2D classification, ab initio model generation, 3D refinements and local resolution estimation were carried out in cryoSPARC 2.14 (Punjani et al., 2017). The 3D reconstructions were performed using C3 symmetry for the RBDs down map while C1 symmetry was applied for the 1 RBD up map. Local refinement (beta) with C1 symmetry was performed on the RBDs down map using a mask focused on the C-terminal helices, which included the bottom of the S2 subunit spanning from residue 1070 to the C terminus. The overall resolution was 3.45 Å for the RBDs down map, 4.00 Å for the 1 RBD up map, and 4.28 Å for the RBDs down local refinement map, confirmed by providing the half maps to Resolve Cryo EM in Phenix (Adams et al., 2004).

#### Cryo-EM model fitting

The coordinates of SARS CoV-2 spike ectodomain structures, PDB entries 6VXX and 6VYB (Walls et al., 2020), were employed as initial models for fitting the cryo-EM map of the RBDs down and the 1 RBD up conformation respectively. In both structures, the RBDs in the “down” position were modeled using the crystallographic structure of RBD in complex with CR3022 Fab (PDB entry 6W41) (Yuan et al., 2020) as a template. For the RBD in the “up” position the map quality did not allow visualization of most side chains which were then modeled as poly-Ala, consistent with what observed for 6VYB (Walls et al., 2020). Manual and automated model building were iteratively performed using Coot (Emsley and Cowtan, 2004) and real space refinement in Phenix respectively, in order to accurately fit the coordinates to the electron density map. Half maps were provided to Resolve Cryo EM tool in Phenix to support manual model building. EMRinger (Barad et al., 2015) and Molprobity (Davis et al., 2004) were used to validate geometry and check structure quality at each iteration step. UCSF Chimera (Pettersen et al., 2004) was used for map-fitting cross correlation calculation (Fit-in-Map tool) and for figure preparation.

#### Probe conjugation

Biotinylated SARS-CoV-2 spike probes were conjugated using either allophycocyanin (APC)-, phycoerythrin (PE)-, BV421-, BV711-, or BV786-labeled streptavidin. Reactions were prepared at a 4:1 molecular ratio of protein to streptavidin, with every monomer labeled. Labeled streptavidin was added in □ increments, with incubations at 4°C (rotating) for 20 minutes in between each addition.

#### Monoclonal yeast display analysis

Yeast display analysis using monoclonal yeast were created, expressed and analyzed as previously published (Wang et al., 2018a). Briefly, VH and VL regions of S652-109, S652-112 and S652-118 were codon optimized for yeast expression, synthesized and cloned into pCT-VHVL-K1 or pCT-VHVL-L1 yeast expression vectors (Genscript). Yeast display vectors were linearized and Saccharomyces cerevisiae strain AWY101 (MATα AGA1::GAL1-AGA1::URA3 PDI1::GAPDH-PDI1::LEU2 ura3-52 trp1 leu2Δ1 his3Δ200 pep4::HIS3 prb1Δ1.6R can1 GAL) was transformed. Yeast cells were maintained in YPD medium (20 g/l dextrose, 20 g/l peptone, and 10 g/l yeast extract); after library transformation, yeast cells were maintained in SDCAA medium (20 g/l dextrose, 6.7 g/l yeast nitrogen base, 5 g/l casamino acids, 8.56 g/l NaH2PO4·H2O, and 10.2 g/l Na2HPO4·7H2O). Fab display was induced by incubating yeast in SGDCAA medium (SDCAA with 20 g/l galactose, 2 g/l dextrose). Two days after induction, 1×106 yeast were incubated in staining buffer (phosphate buffered saline + 0.5% BSA + 2mM EDTA) containing anti-Flag fluorescein isothiocyanate antibody (2 µg/mL; clone M2, Sigma-Aldrich) and the probes for 30 minutes on ice prior to washing 2 times in ice cold staining buffer. Fab expressing yeast (FLAG+) were analyzed for their capacity to bind to the indicated probes using a FACS Aria II (BD Biosciences). For single probe stain experiments yeast were incubated with 4 ng/µl of S2P (APC) for mAb S652-109 and 2 ng/µl for mAbs S652-112 and S652-118, 2 ng/ul of RBD-SD1 (BV421) or 2 ng/µl of RBD-L455RA475RG520R (BV786). Dual stain experiments used 4 ng/µl of S2P (APC), 2 ng/µl of RBD-SD1 (BV421) or 2 ng/µl of RBD-L455RA475RG520R (BV786).

#### Experimental sorting of natively paired antibody heavy:light yeast display libraries

Non-B cells were depleted (EasySep Human B cell enrichment kit w/o CD43 Depletion, STEMCELL Technologies, Vancouver, Canada) and CD27^+^ antigen-experienced B cells were isolated by positive magnetic bead separation (CD27 Human Microbeads, Miltenyi Biotec, Auburn, CA, respectively) (Lagerman et al., 2019). Antigen-experienced B cells were stimulated *in vitro* for five days to enhance antibody gene transcription in the presence of Iscove’s Modified Dulbecco’s Medium (IMDM) (ThermoFisher Scientific, NY, USA) supplemented with 10% FBS, 1 × GlutaMAX, 1 × non-essential amino acids, 1 × sodium pyruvate and 1 × penicillin/streptomycin (Life Technologies, Carlsbad, California, USA) along with 100 units/mL IL-2 and 50 ng/mL IL-21 (PeproTech, Rocky Hill, NJ, USA), and were co-cultured with irradiated 3T3-CD40L fibroblast cells that secrete CD40L to aid B cell expansion. Stimulated B cells were emulsified in the presence of lysis buffer and magnetic beads for mRNA capture as previously described (DeKosky et al., 2015). Magnetic beads were collected and re-emulsified in an overlap-extension RT-PCR emulsion (SuperScript™ III One-Step RT-PCR System with Platinum™ Taq DNA Polymerase, ThermoFisher Scientific, NY, USA) to generate linked VH:VL amplicons (DeKosky et al., 2013; DeKosky et al., 2015; McDaniel et al., 2016; Wang et al., 2018a). cDNA was extracted and a nested PCR was performed (Kapa HiFi HotStart PCR Kit, Kapa Biosystems) to generate ∼850-bp VH:VL products for library cloning into yeast display. 100 ng of natively paired cDNA was amplified with primers containing Not1 and AscI restriction sites for cloning into bidirectional yeast display plasmids (Wang et al., 2018a). Libraries were transformed for amplification in *E. coli*, followed by plasmid DNA extraction and subcloning of a galactose-inducible bidirectional promoter. The resulting native Fab display libraries were co-transformed into electrocompetent AWY101 with AscI/NotI digested pCT vector into yeast cells, and passaged twice before screening (Wang et al., 2018a). Yeast libraries were cultured in SGDCAA medium to induce Fab surface-expression at 20°C and 225 rpm for 36 hours prior to antigen staining. Libraries were stained with an anti-FLAG-FITC monoclonal antibody (clone M2, Sigma-Aldrich, St. Louis, MO), and either 20nM of fluorescently labeled S2P or 100nM of fluorescently labeled NTD or RBD probes, respectively, to isolate antigen binding Fabs. Yeast cells that were gated for Fab expression and antigen binding and collected via fluorescence-activated cell sorting (FACS) and cultured for 48 hours. Libraries were re-induced and re-stained for additional rounds of selection and for the analysis of antigen binding after sorting.

#### PBMC B cell stain

Frozen PBMCs were thawed and immediately stained for viability using the Fixable Aqua Dead Cell Stain Kit (Thermofisher). The PBMCs were stained with the following human surface markers: IgG (G18-145), IgA (S11-8E10), IgM (G20-127), CD8 (RPA-T8), CD3 (OKT3), CD56 (HCD56), CD14 (M5E2), CD27 (O323), CD19 (J3-119), CD38 (HIT2), and CD20 (2H7), sourced from BD, Biolegend, Beckman, and Miltenyi, as well as SARS-CoV-2 probes (2019 S-2P, RBD, RBD-SD-1, RBD ACE2KO, and NTD) in Brilliant Stain Buffer (BD). Samples were collected on a BD FACS-ARIA III and analyzed with FlowJo v10.6.1.

### QUANTIFICATION AND STATISTICAL ANALYSIS

The statistical analyses for the BLI assessment of probe-antibody binding were performed using GraphPad Prism. The SPR data were processed and fit to a 1:1 binding model using Scrubber 2.0 (BioLogic Software).

## References

Adams, P.D., Gopal, K., Grosse-Kunstleve, R.W., Hung, L.W., Ioerger, T.R., McCoy, A.J., Moriarty, N.W., Pai, R.K., Read, R.J., Romo, T.D., et al. (2004). Recent developments in the PHENIX software for automated crystallographic structure determination. J Synchrotron Radiat 11, 53–55.

Antoniou, G., Papakyriacou, I., and Papaneophytou, C. (2017). Optimization of Soluble Expression and Purification of Recombinant Human Rhinovirus Type-14 3C Protease Using Statistically Designed Experiments: Isolation and Characterization of the Enzyme. Mol Biotechnol 59, 407–424.

Barad, B.A., Echols, N., Wang, R.Y., Cheng, Y., DiMaio, F., Adams, P.D., and Fraser, J.S. (2015). EMRinger: side chain-directed model and map validation for 3D cryo-electron microscopy. Nat Methods 12, 943–946.

Brouwer, P.J.M., Caniels, T.G., van der Straten, K., Snitselaar, J.L., Aldon, Y., Bangaru, S., Torres, J.L., Okba, N.M.A., Claireaux, M., Kerster, G., et al. (2020). Potent neutralizing antibodies from COVID-19 patients define multiple targets of vulnerability. bioRxiv, 2020.2005.2012.088716.

Callaway, E. (2020). The race for coronavirus vaccines: a graphical guide. Nature 580, 576–577.

Callaway, E., Cyranoski, D., Mallapaty, S., Stoye, E., and Tollefson, J. (2020). The coronavirus pandemic in five powerful charts. Nature 579, 482–483.

Cao, Y., Su, B., Guo, X., Sun, W., Deng, Y., Bao, L., Zhu, Q., Zhang, X., Zheng, Y., Geng, C., et al. (2020). Potent neutralizing antibodies against SARS-CoV-2 identified by high-throughput single-cell sequencing of convalescent patients’ B cells. Cell.

Casadevall, A., Dadachova, E., and Pirofski, L.A. (2004). Passive antibody therapy for infectious diseases. Nat Rev Microbiol 2, 695–703.

Chen, X., Li, R., Pan, Z., Qian, C., Yang, Y., You, R., Zhao, J., Liu, P., Gao, L., Li, Z., et al. (2020). Human monoclonal antibodies block the binding of SARS-CoV-2 spike protein to angiotensin converting enzyme 2 receptor. Cell Mol Immunol 17, 647–649.

Chi, X., Yan, R., Zhang, J., Zhang, G., Zhang, Y., Hao, M., Zhang, Z., Fan, P., Dong, Y., Yang, Y., et al. (2020). A potent neutralizing human antibody reveals the N-terminal domain of the Spike protein of SARS-CoV-2 as a site of vulnerability. bioRxiv, 2020.2005.2008.083964.

Cordingley, M.G., Callahan, P.L., Sardana, V.V., Garsky, V.M., and Colonno, R.J. (1990). Substrate requirements of human rhinovirus 3C protease for peptide cleavage in vitro. J Biol Chem 265, 9062–9065.

Corti, D., Misasi, J., Mulangu, S., Stanley, D.A., Kanekiyo, M., Wollen, S., Ploquin, A., Doria-Rose, N.A., Staupe, R.P., Bailey, M., et al. (2016). Protective monotherapy against lethal Ebola virus infection by a potently neutralizing antibody. Science 351, 1339–1342.

Cucinotta, D., and Vanelli, M. (2020). WHO Declares COVID-19 a Pandemic. Acta Biomed 91, 157–160.

Davis, I.W., Murray, L.W., Richardson, J.S., and Richardson, D.C. (2004). MOLPROBITY: structure validation and all-atom contact analysis for nucleic acids and their complexes. Nucleic Acids Res 32, W615–619.

DeKosky, B.J., Ippolito, G.C., Deschner, R.P., Lavinder, J.J., Wine, Y., Rawlings, B.M., Varadarajan, N., Giesecke, C., Dorner, T., Andrews, S.F., et al. (2013). High-throughput sequencing of the paired human immunoglobulin heavy and light chain repertoire. Nat Biotechnol 31, 166–169.

DeKosky, B.J., Kojima, T., Rodin, A., Charab, W., Ippolito, G.C., Ellington, A.D., and Georgiou, G. (2015). In-depth determination and analysis of the human paired heavy- and light-chain antibody repertoire. Nat Med 21, 86–91.

Efimov, V.P., Nepluev, I.V., Sobolev, B.N., Zurabishvili, T.G., Schulthess, T., Lustig, A., Engel, J., Haener, M., Aebi, U., Venyaminov, S., and et al. (1994). Fibritin encoded by bacteriophage T4 gene wac has a parallel triple-stranded alpha-helical coiled-coil structure. J Mol Biol 242, 470–486.

Emsley, P., and Cowtan, K. (2004). Coot: model-building tools for molecular graphics. Acta Crystallogr D Biol Crystallogr 60, 2126–2132.

Gaudinski, M.R., Coates, E.E., Novik, L., Widge, A., Houser, K.V., Burch, E., Holman, L.A., Gordon, I.J., Chen, G.L., Carter, C., et al. (2019). Safety, tolerability, pharmacokinetics, and immunogenicity of the therapeutic monoclonal antibody mAb114 targeting Ebola virus glycoprotein (VRC 608): an open-label phase 1 study. Lancet 393, 889–898.

Graham, B.S., and Ambrosino, D.M. (2015). History of passive antibody administration for prevention and treatment of infectious diseases. Curr Opin HIV AIDS 10, 129–134.

Graham, R.L., Donaldson, E.F., and Baric, R.S. (2013). A decade after SARS: strategies for controlling emerging coronaviruses. Nat Rev Microbiol 11, 836–848.

Hoffmann, M., Kleine-Weber, H., Schroeder, S., Kruger, N., Herrler, T., Erichsen, S., Schiergens, T.S., Herrler, G., Wu, N.H., Nitsche, A., et al. (2020). SARS-CoV-2 Cell Entry Depends on ACE2 and TMPRSS2 and Is Blocked by a Clinically Proven Protease Inhibitor. Cell 181, 271–280 e278.

Jager, V., Bussow, K., Wagner, A., Weber, S., Hust, M., Frenzel, A., and Schirrmann, T. (2013). High level transient production of recombinant antibodies and antibody fusion proteins in HEK293 cells. BMC Biotechnol 13, 52.

Jiang, S., Hillyer, C., and Du, L. (2020). Neutralizing Antibodies against SARS-CoV-2 and Other Human Coronaviruses. Trends Immunol 41, 355–359.

Johnson, G., and Wu, T.T. (2000). Kabat database and its applications: 30 years after the first variability plot. Nucleic Acids Res 28, 214–218.

Joyce, M.G., Wheatley, A.K., Thomas, P.V., Chuang, G.Y., Soto, C., Bailer, R.T., Druz, A., Georgiev, I.S., Gillespie, R.A., Kanekiyo, M., et al. (2016). Vaccine-Induced Antibodies that Neutralize Group 1 and Group 2 Influenza A Viruses. Cell 166, 609–623.

Ju, B., Zhang, Q., Ge, J., Wang, R., Sun, J., Ge, X., Yu, J., Shan, S., Zhou, B., Song, S., et al. (2020). Human neutralizing antibodies elicited by SARS-CoV-2 infection. Nature.

Korber, B., Fischer, W., Gnanakaran, S., Yoon, H., Theiler, J., Abfalterer, W., Foley, B., Giorgi, E., Bhattacharya, T., Parker, M., et al. (2020). Spike mutation pipeline reveals the emergence of a more transmissible form of SARS-CoV-2. bioRxiv, 2020.2004.2029.069054.

Lagerman, C.E., Lopez Acevedo, S.N., Fahad, A.S., Hailemariam, A.T., Madan, B., and DeKosky, B.J. (2019). Ultrasonically-guided flow focusing generates precise emulsion droplets for high-throughput single cell analyses. J Biosci Bioeng 128, 226–233.

Lan, J., Ge, J., Yu, J., Shan, S., Zhou, H., Fan, S., Zhang, Q., Shi, X., Wang, Q., Zhang, L., and Wang, X. (2020). Structure of the SARS-CoV-2 spike receptor-binding domain bound to the ACE2 receptor. Nature 581, 215–220.

Liu, L., Xie, J., Sun, J., Han, Y., Zhang, C., Fan, H., Liu, Z., Qiu, Z., He, Y., and Li, T. (2011). Longitudinal profiles of immunoglobulin G antibodies against severe acute respiratory syndrome coronavirus components and neutralizing activities in recovered patients. Scand J Infect Dis 43, 515–521.

Liu, X., Gao, F., Gou, L., Chen, Y., Gu, Y., Ao, L., Shen, H., Hu, Z., Guo, X., and Gao, W. (2020). Neutralizing Antibodies Isolated by a site-directed Screening have Potent Protection on SARS-CoV-2 Infection. bioRxiv, 2020.2005.2003.074914.

Mastronarde, D.N. (2005). Automated electron microscope tomography using robust prediction of specimen movements. J Struct Biol 152, 36–51.

McDaniel, J.R., DeKosky, B.J., Tanno, H., Ellington, A.D., and Georgiou, G. (2016). Ultra-high-throughput sequencing of the immune receptor repertoire from millions of lymphocytes. Nat Protoc 11, 429–442.

Miroshnikov, K.A., Marusich, E.I., Cerritelli, M.E., Cheng, N., Hyde, C.C., Steven, A.C., and Mesyanzhinov, V.V. (1998). Engineering trimeric fibrous proteins based on bacteriophage T4 adhesins. Protein Eng 11, 329–332.

Misasi, J., Gilman, M.S., Kanekiyo, M., Gui, M., Cagigi, A., Mulangu, S., Corti, D., Ledgerwood, J.E., Lanzavecchia, A., Cunningham, J., et al. (2016). Structural and molecular basis for Ebola virus neutralization by protective human antibodies. Science 351, 1343–1346.

Ou, X., Liu, Y., Lei, X., Li, P., Mi, D., Ren, L., Guo, L., Guo, R., Chen, T., Hu, J., et al. (2020). Characterization of spike glycoprotein of SARS-CoV-2 on virus entry and its immune cross-reactivity with SARS-CoV. Nat Commun 11, 1620.

Pallesen, J., Wang, N., Corbett, K.S., Wrapp, D., Kirchdoerfer, R.N., Turner, H.L., Cottrell, C.A., Becker, M.M., Wang, L., Shi, W., et al. (2017). Immunogenicity and structures of a rationally designed prefusion MERS-CoV spike antigen. Proc Natl Acad Sci U S A 114, E7348–E7357.

Perera, R.A., Mok, C.K., Tsang, O.T., Lv, H., Ko, R.L., Wu, N.C., Yuan, M., Leung, W.S., Chan, J.M., Chik, T.S., et al. (2020). Serological assays for severe acute respiratory syndrome coronavirus 2 (SARS-CoV-2), March 2020. Euro Surveill 25.

Pettersen, E.F., Goddard, T.D., Huang, C.C., Couch, G.S., Greenblatt, D.M., Meng, E.C., and Ferrin, T.E. (2004). UCSF Chimera--a visualization system for exploratory research and analysis. J Comput Chem 25, 1605–1612.

Pinto, D., Park, Y.-J., Beltramello, M., Walls, A.C., Tortorici, M.A., Bianchi, S., Jaconi, S., Culap, K., Zatta, F., De Marco, A., et al. (2020). Structural and functional analysis of a potent sarbecovirus neutralizing antibody. bioRxiv, 2020.2004.2007.023903.

Punjani, A., Rubinstein, J.L., Fleet, D.J., and Brubaker, M.A. (2017). cryoSPARC: algorithms for rapid unsupervised cryo-EM structure determination. Nat Methods 14, 290–296.

Ridgway, J.B., Presta, L.G., and Carter, P. (1996). ’Knobs-into-holes’ engineering of antibody CH3 domains for heavy chain heterodimerization. Protein Eng 9, 617–621.

Robbiani, D.F., Gaebler, C., Muecksch, F., Lorenzi, J.C.C., Wang, Z., Cho, A., Agudelo, M., Barnes, C.O., Gazumyan, A., Finkin, S., et al. (2020). Convergent Antibody Responses to SARS-CoV-2 Infection in Convalescent Individuals. bioRxiv, 2020.2005.2013.092619.

Rogers, T.F., Zhao, F., Huang, D., Beutler, N., Burns, A., He, W.-t., Limbo, O., Smith, C., Song, G., Woehl, J., et al. (2020). Rapid isolation of potent SARS-CoV-2 neutralizing antibodies and protection in a small animal model. bioRxiv, 2020.2005.2011.088674.

Scheres, S.H. (2012). RELION: implementation of a Bayesian approach to cryo-EM structure determination. J Struct Biol 180, 519–530.

Seydoux, E., Homad, L.J., MacCamy, A.J., Parks, K.R., Hurlburt, N.K., Jennewein, M.F., Akins, N.R., Stuart, A.B., Wan, Y.-H., Feng, J., et al. (2020). Characterization of neutralizing antibodies from a SARS-CoV-2 infected individual. bioRxiv, 2020.2005.2012.091298.

Shen, Y., Maupetit, J., Derreumaux, P., and Tufféry, P. (2014). Improved PEP-FOLD Approach for Peptide and Miniprotein Structure Prediction. Journal of Chemical Theory and Computation 10, 4745–4758.

Suloway, C., Pulokas, J., Fellmann, D., Cheng, A., Guerra, F., Quispe, J., Stagg, S., Potter, C.S., and Carragher, B. (2005). Automated molecular microscopy: the new Leginon system. J Struct Biol 151, 41–60.

Thevenet, P., Shen, Y., Maupetit, J., Guyon, F., Derreumaux, P., and Tuffery, P. (2012). PEP-FOLD: an updated de novo structure prediction server for both linear and disulfide bonded cyclic peptides. Nucleic Acids Res 40, W288–293.

Tian, X., Li, C., Huang, A., Xia, S., Lu, S., Shi, Z., Lu, L., Jiang, S., Yang, Z., Wu, Y., and Ying, T. (2020). Potent binding of 2019 novel coronavirus spike protein by a SARS coronavirus-specific human monoclonal antibody. Emerg Microbes Infect 9, 382–385.

Walker, L.M., and Burton, D.R. (2018). Passive immunotherapy of viral infections: ‘super-antibodies’ enter the fray. Nat Rev Immunol 18, 297–308.

Walls, A.C., Park, Y.J., Tortorici, M.A., Wall, A., McGuire, A.T., and Veesler, D. (2020). Structure, Function, and Antigenicity of the SARS-CoV-2 Spike Glycoprotein. Cell 181, 281–292 e286.

Wang, B., DeKosky, B.J., Timm, M.R., Lee, J., Normandin, E., Misasi, J., Kong, R., McDaniel, J.R., Delidakis, G., Leigh, K.E., et al. (2018a). Functional interrogation and mining of natively paired human VH:VL antibody repertoires. Nat Biotechnol 36, 152–155.

Wang, C., Li, W., Drabek, D., Okba, N.M.A., van Haperen, R., Osterhaus, A., van Kuppeveld, F.J.M., Haagmans, B.L., Grosveld, F., and Bosch, B.J. (2020a). A human monoclonal antibody blocking SARS-CoV-2 infection. Nat Commun 11, 2251.

Wang, L., Shi, W., Chappell, J.D., Joyce, M.G., Zhang, Y., Kanekiyo, M., Becker, M.M., van Doremalen, N., Fischer, R., Wang, N., et al. (2018b). Importance of Neutralizing Monoclonal Antibodies Targeting Multiple Antigenic Sites on the Middle East Respiratory Syndrome Coronavirus Spike Glycoprotein To Avoid Neutralization Escape. J Virol 92.

Wang, Q., Zhang, Y., Wu, L., Niu, S., Song, C., Zhang, Z., Lu, G., Qiao, C., Hu, Y., Yuen, K.Y., et al. (2020b). Structural and Functional Basis of SARS-CoV-2 Entry by Using Human ACE2. Cell 181, 894–904 e899.

Whittle, J.R., Wheatley, A.K., Wu, L., Lingwood, D., Kanekiyo, M., Ma, S.S., Narpala, S.R., Yassine, H.M., Frank, G.M., Yewdell, J.W., et al. (2014). Flow cytometry reveals that H5N1 vaccination elicits cross-reactive stem-directed antibodies from multiple Ig heavy-chain lineages. J Virol 88, 4047–4057.

Wrapp, D., De Vlieger, D., Corbett, K.S., Torres, G.M., Wang, N., Van Breedam, W., Roose, K., van Schie, L., Team, V.-C.C.-R., Hoffmann, M., et al. (2020a). Structural Basis for Potent Neutralization of Betacoronaviruses by Single-Domain Camelid Antibodies. Cell 181, 1004–1015 e1015.

Wrapp, D., Wang, N., Corbett, K.S., Goldsmith, J.A., Hsieh, C.L., Abiona, O., Graham, B.S., and McLellan, J.S. (2020b). Cryo-EM structure of the 2019-nCoV spike in the prefusion conformation. Science 367, 1260–1263.

Wu, X., Yang, Z.Y., Li, Y., Hogerkorp, C.M., Schief, W.R., Seaman, M.S., Zhou, T., Schmidt, S.D., Wu, L., Xu, L., et al. (2010). Rational design of envelope identifies broadly neutralizing human monoclonal antibodies to HIV-1. Science 329, 856–861.

Wu, X., Zhou, T., Zhu, J., Zhang, B., Georgiev, I., Wang, C., Chen, X., Longo, N.S., Louder, M., McKee, K., et al. (2011). Focused evolution of HIV-1 neutralizing antibodies revealed by structures and deep sequencing. Science 333, 1593–1602.

Wu, Y., Wang, F., Shen, C., Peng, W., Li, D., Zhao, C., Li, Z., Li, S., Bi, Y., Yang, Y., et al. (2020). A noncompeting pair of human neutralizing antibodies block COVID-19 virus binding to its receptor ACE2. Science 368, 1274–1278.

Yan, R., Zhang, Y., Li, Y., Xia, L., Guo, Y., and Zhou, Q. (2020a). Structural basis for the recognition of SARS-CoV-2 by full-length human ACE2. Science 367, 1444–1448.

Yan, Y., Chang, L., and Wang, L. (2020b). Laboratory testing of SARS-CoV, MERS-CoV, and SARS-CoV-2 (2019-nCoV): Current status, challenges, and countermeasures. Rev Med Virol 30, e2106.

Yongchen, Z., Shen, H., Wang, X., Shi, X., Li, Y., Yan, J., Chen, Y., and Gu, B. (2020). Different longitudinal patterns of nucleic acid and serology testing results based on disease severity of COVID-19 patients. Emerg Microbes Infect 9, 833–836.

Yuan, M., Wu, N.C., Zhu, X., Lee, C.D., So, R.T.Y., Lv, H., Mok, C.K.P., and Wilson, I.A. (2020). A highly conserved cryptic epitope in the receptor binding domains of SARS-CoV-2 and SARS-CoV. Science 368, 630–633.

Zeng, X., Li, L., Lin, J., Li, X., Liu, B., Kong, Y., Zeng, S., Du, J., Xiao, H., Zhang, T., et al. (2020). Isolation of a human monoclonal antibody specific for the receptor binding domain of SARS-CoV-2 using a competitive phage biopanning strategy. Antib Ther, tbaa008.

Zhao, R., Li, M., Song, H., Chen, J., Ren, W., Feng, Y., Gao, G.F., Song, J., Peng, Y., Su, B., et al. (2020). Early detection of SARS-CoV-2 antibodies in COVID-19 patients as a serologic marker of infection. Clin Infect Dis, Epub Date 2020/0502, PMCID PMC7197602.

Zhou, G., and Zhao, Q. (2020). Perspectives on therapeutic neutralizing antibodies against the Novel Coronavirus SARS-CoV-2. Int J Biol Sci 16, 1718–1723.

Zost, S.J., Gilchuk, P., Chen, R.E., Case, J.B., Reidy, J.X., Trivette, A., Nargi, R.S., Sutton, R.E., Suryadevara, N., Chen, E.C., et al. (2020). Rapid isolation and profiling of a diverse panel of human monoclonal antibodies targeting the SARS-CoV-2 spike protein. bioRxiv, 2020.2005.2012.091462.

