## Supplemental Information for "Structure-Based Design with Tag-Based Purification and In-Process Biotinylation Enable Streamlined Development of SARS-CoV-2 Spike Molecular Probes"

**Table S1. Plasmid Sequences of SARS-CoV-2 Molecular Probes, Related to Figures 2, 4, and 6.**

See Excel spreadsheet file TableS1.xlsx

**Table S2. Cryo-EM Data Collection and Refinement Statistics, Related to Figures 3 and 4.**

| | Biotinylated SARS-CoV2-S2P-AVI probe (RBDs down) | Biotinylated SARS-CoV2-S2P-AVI probe (1 RBD up) |
|---|---|---|
| **EMDB ID** | EMD-22161 | EMD-22162 |
| **PDB ID** | 6XF5 | 6XF6 |
| Data Collection | | |
| Microscope | FEI Titan Krios | FEI Titan Krios |
| Voltage (kV) | 300 | 300 |
| Electron dose ($e^-/Å^2$) | 41.92 | 41.92 |
| Detector | Gatan K3 BioQuantum | Gatan K3 BioQuantum |
| Pixel Size (Å) | 1.07 | 1.07 |
| Defocus Range (µm) | -0.8/-2.5 | -0.8/-2.5 |
| Magnification | 81000 | 81000 |
| Reconstruction | | |
| Software | cryoSPARC v2.14 | cryoSPARC v2.14 |
| Particles | 43,176 | 34,223 |
| Symmetry | C3 | C1 |
| Box size (pix) | 440 | 440 |
| Resolution (Å) ($FSC_{0.143}$) | 3.45 | 4.00 |
| Refinement | | |
| Software | Phenix 1.18 | Phenix 1.18 |
| Protein residues | 3027 | 2928 |
| Chimera CC | 0.82 | 0.86 |
| EMRinger Score | 1.58 | 1.22 |
| R.m.s. deviations | | |
| Bond lengths (Å) | 0.004 | 0.003 |
| Bond angles (°) | 0.778 | 0.689 |
| Validation | | |
| Molprobity score | 1.53 | 1.28 |
| Clash score | 4.94 | 3.90 |
| Favored rotamers (%) | 98.7 | 100 |
| Ramachandran | | |
| Favored regions (%) | 96.8 | 97.5 |
| Allowed regions (%) | 3.2 | 2.5 |
| Disallowed regions (%) | 0 | 0 |

### Figure S1

**A**

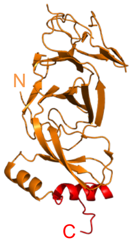


```
MGWSCIILFLVATATGVHSAPELLGGPSVFLFPPKPKDTLMISRTPEVTCVVVDVSHEDPE
VKFNWYVDGVEVHNAKTKPREEQYNSTYRVVSVLTVLHQDWLNGKEYKCKVSNKALPAPIE
KTISKAKGQPREPQVYTLPPSRDELTKNQVSLYCLVKGFYPSDIAVEWESNGQPENNYKTT
PPVLDSDGSFFLYSKLTVDKSRWQQGNVFSCSVMHEALHNHYTQKSLSLSPGKGGGGSGGG
GSGGGGSGGGGSAPELLGGPSVFLFPPKPKDTLMISRTPEVTCVVVDVSHEDPEVKFNWYV
DGVEVHNAKTKPREEQYNSTYRVVSVLTVLHQDWLNGKEYKCKVSNKALPAPIEKTISKAK
GQPREPQVYTLPPSRDELTKNQVSLTCLVKGFYPSDIAVEWESNGQPENNYKTTPPVLDSD
GSFFLTSKLTVDKSRWQQGNVFSCSVMHEALHNHYTQKSLSLSPGKGGSGGGGSGGLEVLF
QGPQCVNLTTRTQLPPAYTNSFTRGVYYPDKVFRSSVLHSTQDLFLPFFSNVTWFHAIHVS
GTNGTKRFDNPVLPFNDGVYFASTEKSNIIRGWIFGTTLDSKTQSLLIVNNATNVVIKVCE
FQFCNDPFLGVYYHKNNKSWMESEFRVYSSANNCTFEYVSQPFLMDLEGKQGNFKNLREFV
FKNIDGYFKIYSKHTPINLVRDLPQGFSALEPLVDLPIGINITRFQTLLALHRSYLTPGDS
SSGWTAGAAAYYVGYLQPRTFLLKYNENGTITDAVDCALDPLSETKCTLKSGGGLVPQQSG
GLNDIFEAQKIEWHEG
```

Key regions:

scFc

HRV3C site

NTD

10lnQQ-AVI

**B**

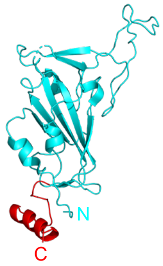


```
MGWSCIILFLVATATGVHSAPELLGGPSVFLFPPKPKDTLMISRTPEVTCVVVDVSHEDPE
VKFNWYVDGVEVHNAKTKPREEQYNSTYRVVSVLTVLHQDWLNGKEYKCKVSNKALPAPIE
KTISKAKGQPREPQVYTLPPSRDELTKNQVSLYCLVKGFYPSDIAVEWESNGQPENNYKTT
PPVLDSDGSFFLYSKLTVDKSRWQQGNVFSCSVMHEALHNHYTQKSLSLSPGKGGGGSGGG
GSGGGGSGGGGSAPELLGGPSVFLFPPKPKDTLMISRTPEVTCVVVDVSHEDPEVKFNWYV
DGVEVHNAKTKPREEQYNSTYRVVSVLTVLHQDWLNGKEYKCKVSNKALPAPIEKTISKAK
GQPREPQVYTLPPSRDELTKNQVSLTCLVKGFYPSDIAVEWESNGQPENNYKTTPPVLDSD
GSFFLTSKLTVDKSRWQQGNVFSCSVMHEALHNHYTQKSLSLSPGKGGSGGGGSGGLEVLF
QGPLCPFGEVFNATRFASVYAWNRKRISNCVADYSVLYNSASFSTFKCYGVSPTKLNDLCF
TNVYADSFVIRGDEVRQIAPGQTGKIADYNYKLPDDFTGCVIAWNSNNLDSKVGGNYNYLY
RLFRKSNLKPFERDISTEIYQAGSTPCNGVEGFNCYFPLQSYGFQPTNGVGYQPYRVVVLS
FELLHAPATVCGGGGLVPQQSGGLNDIFEAQKIEWHEG
```

Key regions:

scFc

HRV3C site

RBD

10lnQQ-AVI

**C**

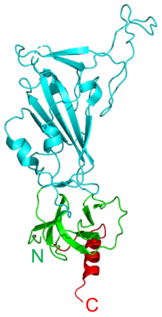


```
MGWSCIILFLVATATGVHSAPELLGGPSVFLFPPKPKDTLMISRTPEVTCVVVDVSHEDPE
VKFNWYVDGVEVHNAKTKPREEQYNSTYRVVSVLTVLHQDWLNGKEYKCKVSNKALPAPIE
KTISKAKGQPREPQVYTLPPSRDELTKNQVSLYCLVKGFYPSDIAVEWESNGQPENNYKTT
PPVLDSDGSFFLYSKLTVDKSRWQQGNVFSCSVMHEALHNHYTQKSLSLSPGKGGGGSGGG
GSGGGGSGGGGSAPELLGGPSVFLFPPKPKDTLMISRTPEVTCVVVDVSHEDPEVKFNWYV
DGVEVHNAKTKPREEQYNSTYRVVSVLTVLHQDWLNGKEYKCKVSNKALPAPIEKTISKAK
GQPREPQVYTLPPSRDELTKNQVSLTCLVKGFYPSDIAVEWESNGQPENNYKTTPPVLDSD
GSFFLTSKLTVDKSRWQQGNVFSCSVMHEALHNHYTQKSLSLSPGKGGSGGGGSGGLEVLF
QGPRVQPTESIVRFPNITNLCPFGEVFNATRFASVYAWNRKRISNCVADYSVLYNSASFST
FKCYGVSPTKLNDLCFTNVYADSFVIRGDEVRQIAPGQTGKIADYNYKLPDDFTGCVIAWN
SNNLDSKVGGNYNYLYRLFRKSNLKPFERDISTEIYQAGSTPCNGVEGFNCYFPLQSYGFQ
PTNGVGYQPYRVVVLSFELLHAPATVCGPKKSTNLVKNKCVNFNFNGLTGTGVLTESNKKF
LPFQQFGRDIADTTDAVRDPQTLEILDITPCSGGGLVPQQSGGLNDIFEAQKIEWHEG
```

Key regions:

scFc

HRV3C site

RBD-SD1

10lnQQ-AVI

**Figure S1. Complete Sequences for Molecular Probes Comprising SARS-CoV-2 NTD, RBD and RBD-SD1 Domains, Related to Figure 4**

(A) Structural model of the NTD probe with the C-terminus 10lnQQ linker and AVI tag highlighted in red (left); the model of the 10lnQQ-AVI was predicted on the PEP-FOLD server. The full sequence of the NTD molecular probe including the N-terminus scFc and HRV3C site and C-terminus 10lnQQ linker and AVI tag is shown to the right.

(B) Structural model of the RBD probe with the C-terminus 10lnQQ linker and AVI tag highlighted in red (left); the model of the 10lnQQ-AVI was predicted on the PEP-FOLD server. The full sequence of the RBD molecular probe including the N-terminus scFc and HRV3C site and C-terminus 10lnQQ linker and AVI tag is shown to the right.

(C) Structural model of the RBD-SD1 probe with the C-terminus 10lnQQ linker and AVI tag highlighted in red (left); the model of the 10lnQQ-AVI was predicted on the PEP-FOLD server. The full sequence of the RBD-SD1 molecular probe including the N-terminus scFc and HRV3C site and C-terminus 10lnQQ linker and AVI tag is shown to the right.

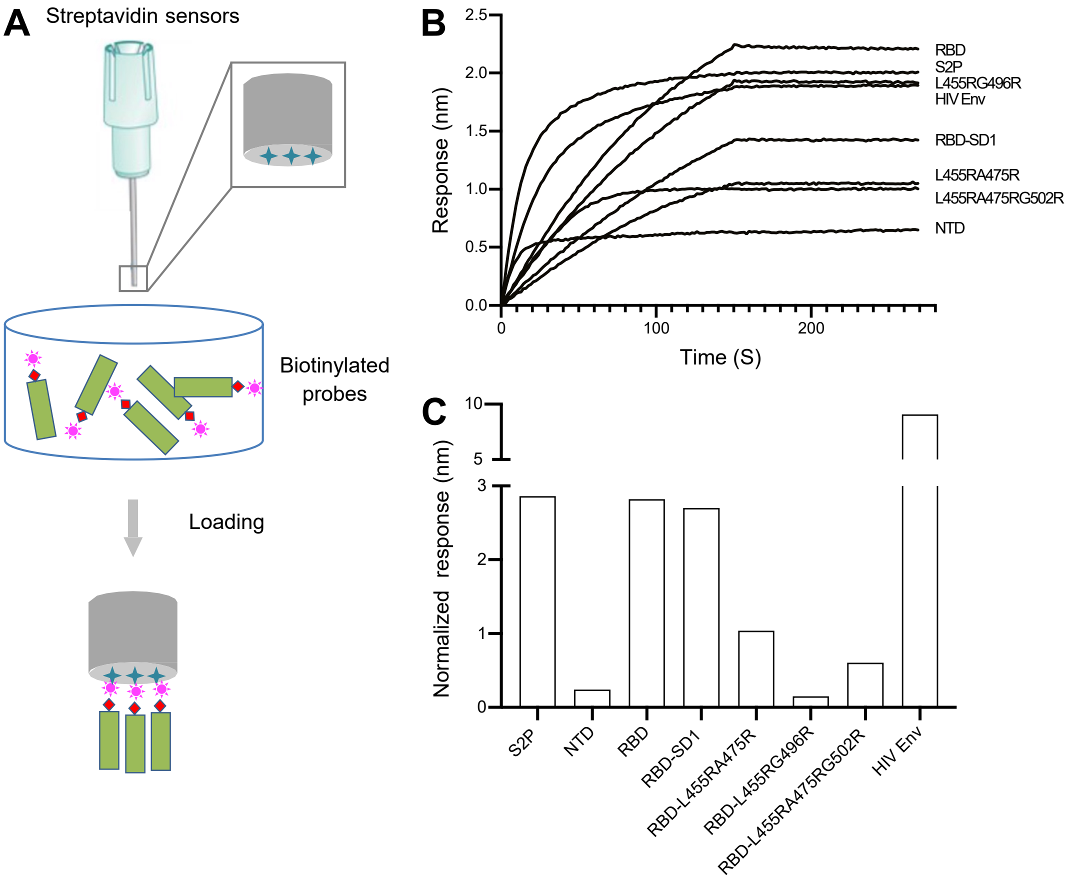


**Figure S2. Assessment Biotinylation Levels of Molecular Probe, Related to Figures 3, 5, and 6**

(A) Schematic of Bio-Layer Interferometry assay. Response of biotinylated molecular probes captured onto the tip of streptavidin sensor was recorded.

(B) Sensorgrams of biotinylated molecular probes binding to streptavidin sensor. Concentrations of probes were adjusted to yield a response >1nm for subsequent antibody or receptor binding studies. Loading concentrations were as follows, S2P (80 μg/ml), HIV Env (20 μg/ml), NTD (40 μg/ml), RBD (5 μg/ml), RBD-SD1 (5 μg/ml), RBD-L455RA475R (7.5 μg/ml), RBD-L455RG496R (80 μg/ml), RBD-L455RA475RG502R (15 μg/ml).

(C) Normalized biotinylation level for different molecular probes at the 200 nM concentration.

### Figure S3

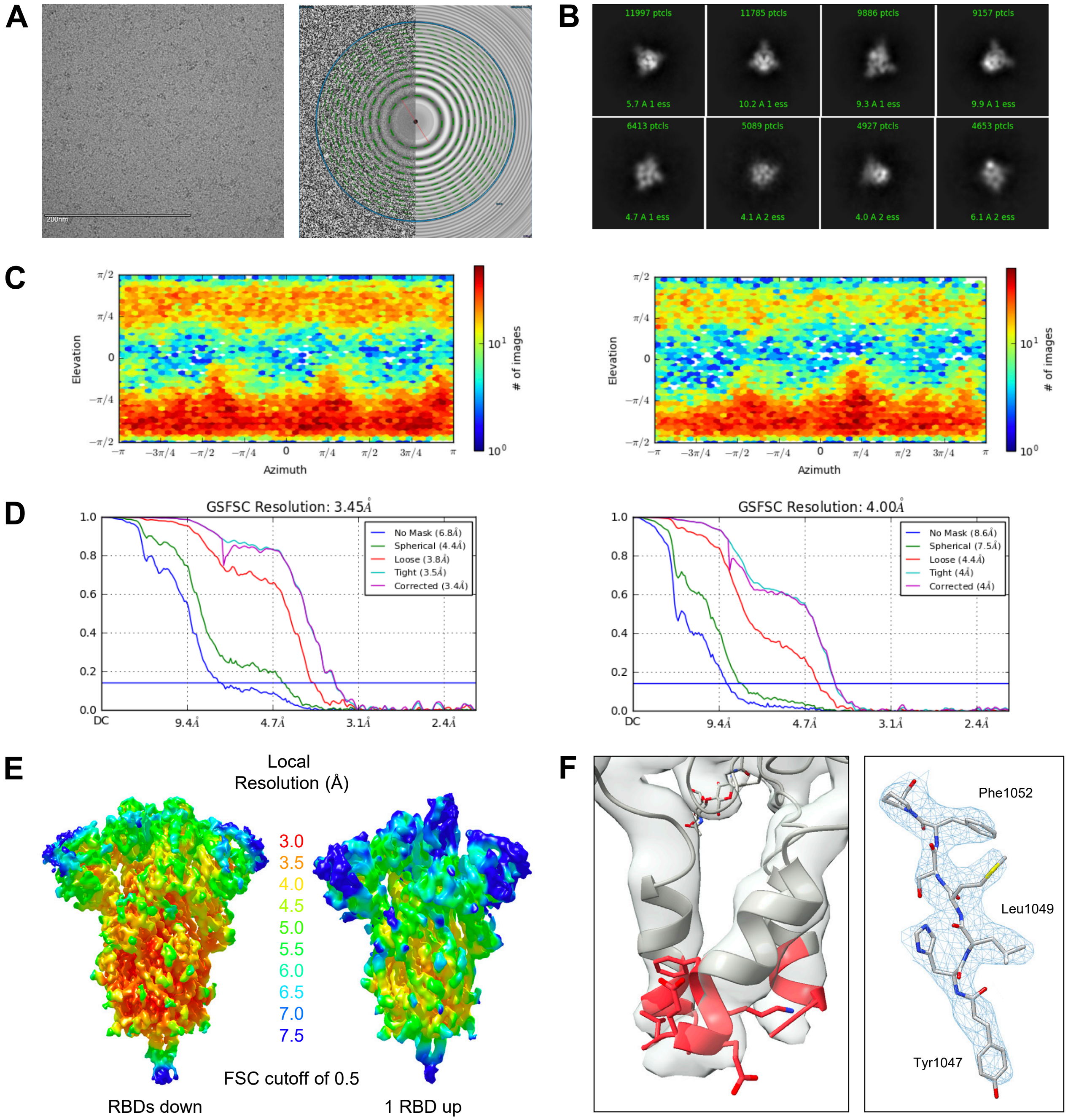


**Figure S3. Cryo-EM Details of Biotinylated SARS-CoV2 S2P Probe, Related to Figures 3 and 4**

(A) Representative micrograph and CTF of the micrograph are shown.

(B) Representative 2D class averages are shown.

(C) The orientations of all particles used in the final refinement are shown as a heatmap (left: RBDs down; right: 1 RBD up).

(D) The gold-standard Fourier shell correlation resulted in a resolution of 3.45 Å using non-uniform refinement with C3 symmetry for the RBDs down state (left) and 4.0 Å using non-uniform refinement with C1 symmetry for the 1 RBD up state (right). The resolution is 4.28 Å for the map resulting from local refinement (beta), obtained using C1 symmetry.

(E) The local resolution of the two maps are shown generated through cryoSPARC using an FSC cutoff of 0.5.

(F) Representative density is shown for the C-terminal helices of the biotinylated SARS-CoV2-S2P-AVI probe. The map resulting from local refinement using C1 symmetry for the RBDs down state shows that the C-terminal helices observed in the previous models (6VSB, 6VYB, 6VXX) include 5 more amino acids (residues 1148-1152) colored in pale red (left panel). Representative cryo-EM density, shown as blue mesh, and model, shown as sticks, for residues 1047-1053 (right panel). Carbon atoms colored in gray, oxygen in red, nitrogen in blue, sulfur in yellow.

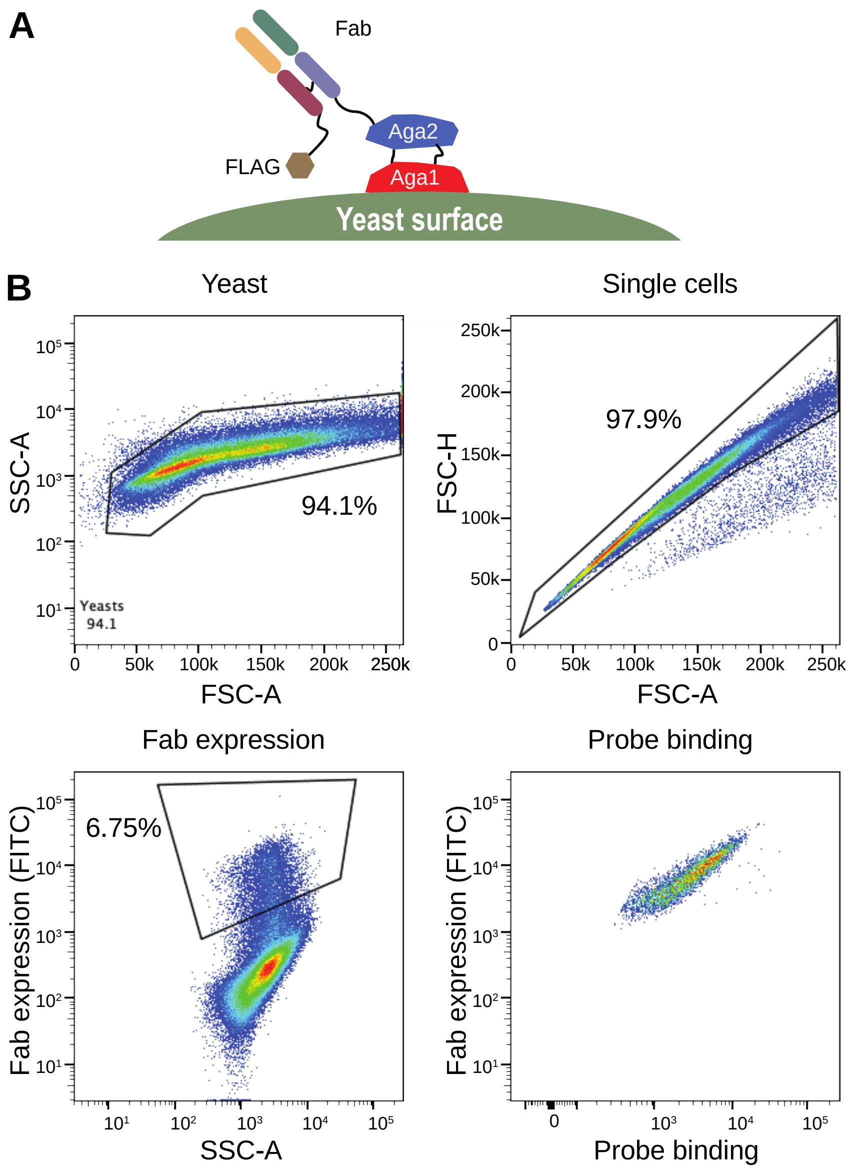


**Figure S4. Yeast Fab Display and Gating Tree for Yeast Display Analysis of Probe Binding, Related to Figure 7**

(A) Saccharomyces cerevisiae strain AWY101 transfected with yeast display vector and Fab display is induced by incubating yeast in galactose containing media. The presence of Fab expressed on the yeast surface can be detected by staining with an anti-Flag antibody and analyzing using flow cytometry.

(B) Induced yeast bearing Fabs of interested are analyzed by the indicated gating strategy. Singlets are analyzed for Fab expression and the proportion of probe binding determined within this population of yeast. Shown is a representative data.

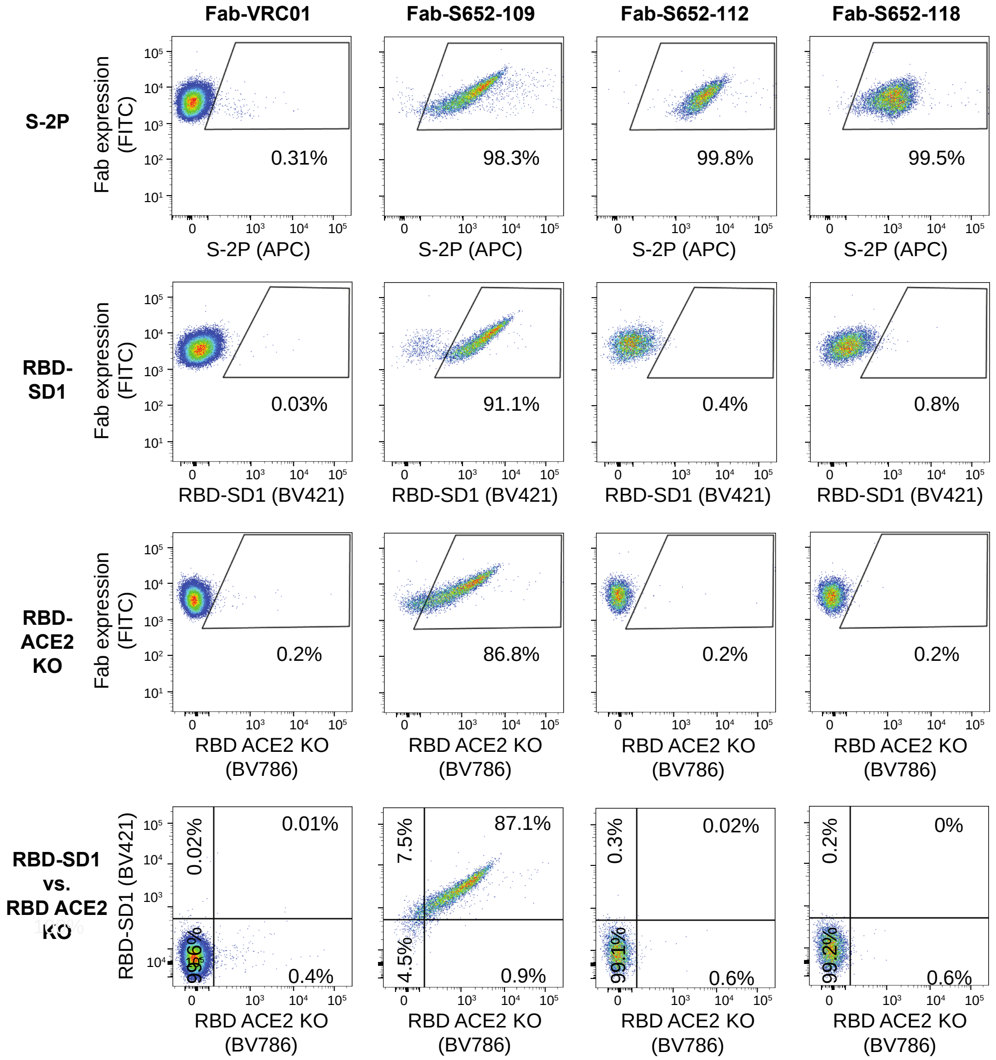


**Figure S5. Yeast SARS-CoV Cross-Reactive Fab Binding to SARS-CoV-2 Antigenic Probes, Related to Figure 7**
Binding of yeast expressing SARS-CoV cross-reactive Fabs (S652-109, S652-112, and S652-118) or HIV targeting VRC01 Fab to SARS-CoV-2 antigenic probes: S2P (APC), RBD-SD1 (BV421) or RBD-ACE2 KO (BV786). RBD-ACE2 KO is the triple-Arg mutant, RBD-L455RA475RG502R.

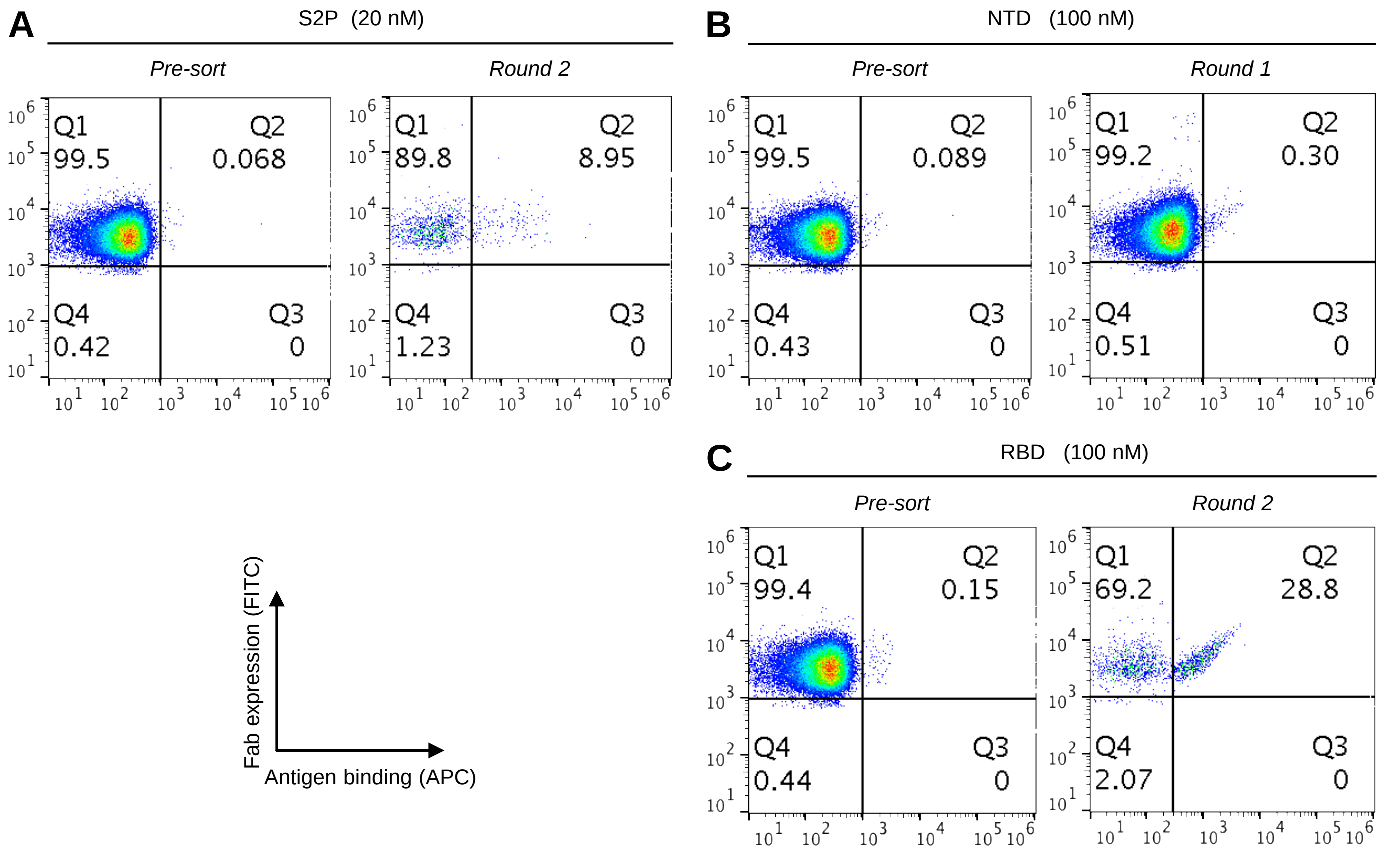


**Figure S6. Natively Paired Heavy:Light Yeast Display Sorting in a Second SARS-CoV-2 Convalescent Donor, Related to Figure 7**

(A) Experimental sorting with the S2P trimer probe at 20 nM concentration.
(B) Experimental sorting with the NTD probe at 100 nM concentration.
(C) Experimental sorting with the RBD probe at 100 nM concentration.

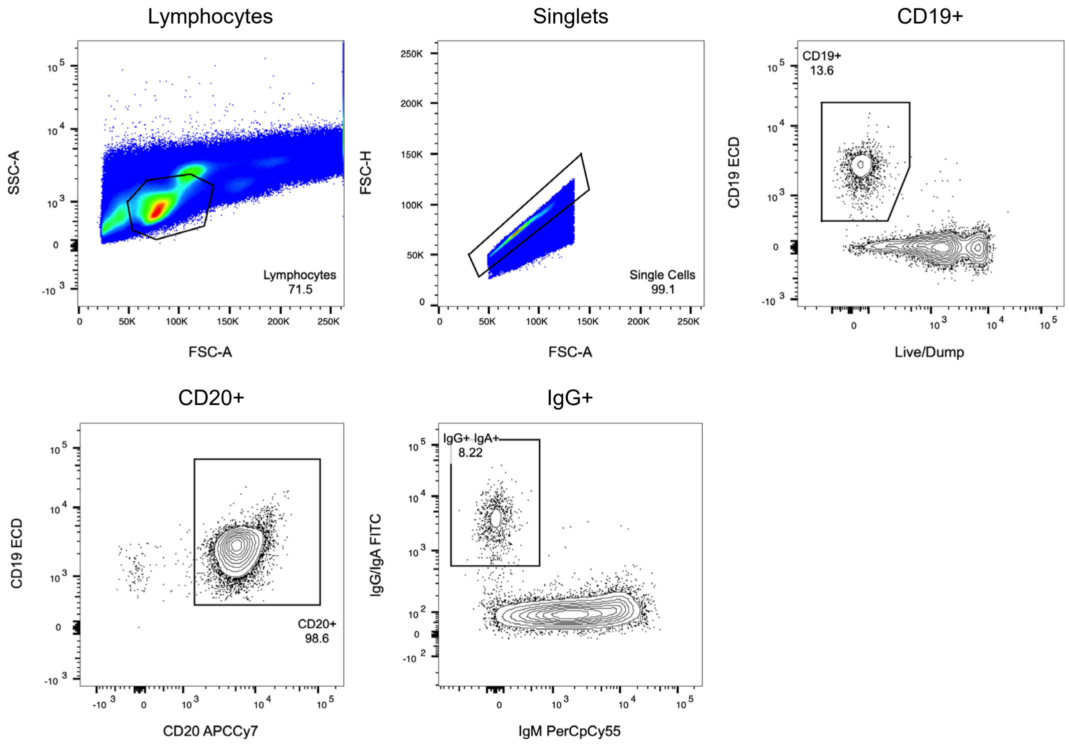


**Figure S7. Flow Cytometry Gating Scheme for Sorting COVID-19 Convalescent PBMC, Relate to Figure 7**
A hierarchical gating strategy was used to identify live, singlet B cells in human PBMC. Bivariate gates for probe-binding B cells presented in Figure 7D are from the IgG+ and IgA+ B cell gate.
